# Differential utilization of PACAP and glutamate in parabrachial neurons shapes pain and stress responses

**DOI:** 10.64898/2026.10.07.755626

**Authors:** Sudhuman Singh, Sunny Zhihong Jiang, William J. Fortún Rosado, William Brancaleone, Diego Castillo-Rolón, Hugo A. Tejeda, Lee E. Eiden, Yarimar Carrasquillo

**Affiliations:** Pain Neurocircuitry and Cellular Plasticity Lab, National Center for Complementary and Integrative Health, National Institute of Mental Health Intramural Research Program, Bethesda, MD, USA 20892; National Institute on Drug Abuse, National Institute of Mental Health Intramural Research Program, Bethesda, 20892; Section on Molecular Neuroscience, National Institute of Mental Health Intramural Research Program, Bethesda, 20892; Section on Neuromodulation and Synaptic Integration, National Institute of Mental Health Intramural Research Program, Bethesda, MD, USA 20892

## Abstract

The external parabrachial nucleus (elPBN) contributes to diverse behaviors, including pain and stress. Neurons in this region use glutamatergic and neuropeptidergic signaling, with pituitary adenylate cyclase-activating polypeptide (PACAP) expressed in a subset of glutamatergic neurons. Deletion of PACAP in elPBN neurons attenuates behavioral responses to stress, but the contribution of PACAP to pain sensitization and of glutamatergic neurotransmission to pain- and stress-related responses are poorly understood. Here, we show that constitutive loss of PACAP markedly attenuated nerve injury-induced hypersensitivity across heat, pressure, cold and tactile modalities without altering baseline nociception, while also reducing inflammation-induced spontaneous pain-related behavior. Selective deletion of PACAP within elPBN neurons phenocopied the suppression of injury-induced hypersensitivity, but left inflammation-induced pain-related behavior unaffected. Ex vivo optogenetic-assisted circuit mapping showed that activation of PACAP-expressing elPBN terminals elicited glutamatergic transmission onto central amygdala (CeA) PKCδ-expressing neurons. Selective deletion of VGluT2 in elPBN neurons reduced nerve injury-induced hypersensitivity but, unlike PACAP deletion, also reduced inflammation-induced pain-related behavior, while leaving stress-induced hypophagia intact. Together, these findings reveal differential requirements for PACAP and glutamatergic neurotransmission within the elPBN to allow pain- and stress-related behaviors.

**Significance Statement:** The parabrachial nucleus contributes to pain and stress responses through neurons that express multiple neurotransmitters. Here, we show that disrupting PACAP or glutamatergic signaling in the elPBN produces distinct effects across nerve injury-induced sensitization, inflammation-induced pain-related behavior, and restraint stress-induced hypophagia. These findings reveal functional specialization of signaling mechanisms within the elPBN, highlighting neurotransmitter diversity as an important feature of its broad behavioral functions.

## Introduction

The external lateral parabrachial nucleus (elPBN) contributed to diverse behavioral responses to pain, stress, and other physiologically important stimuli (Chiang et al., 2019; Palmiter, 2018). This functional diversity is accompanied by substantial cellular heterogeneity within the PBN (Pauli et al., 2022). Single-cell transcriptomic and anatomical studies have identified numerous distinct PBN populations with different distributions and projection patterns. These can support distinct but overlapping functions: for example, Tac1-expressing lateral PBN neurons contribute to nocifensive escape behaviors (Barik et al., 2018), whereas Oprm1-expressing neurons comprise anatomically and functionally distinct core and shell populations that differentially contribute to pain, anxiety, and breathing (Liu et al., 2022). Recent work further shows that sustained pain engages a molecularly distributed population of Y1R-expressing PBN neurons that spans previously defined cell types and is regulated by competing physiological states (Goldstein et al., 2025). Even within pain-related PBN circuits, nerve injury produces distinct changes in excitability across neuronal populations (Torres-Rodriguez et al., 2024). Moreover, many elPBN neurons express both fast neurotransmitters and neuropeptides, and recent studies demonstrate that these signaling molecules can make different contributions to behavioral output (Kim et al., 2024). Together, these findings suggest that the diversity of response patterns of the PBN contribute to behavioral diversity and may arise not only from which neurons are recruited and where they project, but also from the signaling molecules they use.

Pituitary adenylate cyclase-activating polypeptide (PACAP) is expressed in a subset of glutamatergic elPBN neurons (Zhang et al., 2021) and has been implicated in both pain- and stress-related behaviors (Jiang et al., 2023; Missig et al., 2017; Missig et al., 2014; Seiglie et al., 2023). Studies using global PACAP knockout (PACAP-KO) mice demonstrated that loss of endogenous PACAP markedly reduces mechanical and thermal hypersensitivity in neuropathic and inflammatory pain models without affecting baseline nociceptive responses (Mabuchi et al., 2004). Focusing on ascending pathways, Missig et al. (2014) identified a peptidergic projection from lateral PBN neurons to the central amygdala (CeA) and showed that delivery of PACAP into the CeA increased mechanical and thermal sensitivity, anxiety-like behavior, and hypothalamic-pituitary-adrenal axis activation in otherwise naive animals (Missig et al., 2014). In a chronic pain model Missig et al. (2017) found increased PACAP signaling along the PBN→CeA pathway and demonstrated that pharmacological disruption of PACAP receptor signaling in the CeA reverses injury-induced tactile allodynia and affective pain-related behaviors (Missig et al., 2017). PACAP signaling in the elPBN has also been implicated in stress: we previously showed that selective PACAP deletion in elPBN neurons attenuates restraint stress-induced hypophagia and CeA c-Fos recruitment (Jiang et al., 2023). More recently, chemogenetic activation of PACAP-expressing neurons of the PBN was shown to increase tactile sensitivity and anxiety-like behaviors (Seiglie et al., 2023). Together these studies implicate PACAP signaling from the PBN in pain- and stress-related behaviors, albeit indirectly through global gene deletion, exogeneous PACAP or receptor-directed pharmacology, or activation of PACAP-expressing neuronal populations. Whether endogenous PACAP contained within elPBN neurons contributes to nerve injury-induced pain sensitization remains unknown.

In addition to neuropeptides, most PBN neurons use glutamate as a fast excitatory neurotransmitter. The PBN provides a strong glutamatergic input to CeA, and extensive electrophysiological studies have shown that excitatory transmission at the PBN→CeA pathway is potentiated across persistent pain states (Kato, 2018; Neugebauer et al., 2020). Pharmacological and genetic studies further demonstrate that glutamate receptor signaling within the CeA contributes to pain sensitization, including through NMDA and metabotropic glutamate receptor-dependent mechanisms (Kolber et al., 2010; Neugebauer V, 2003). Neuropeptide signaling can also modulate glutamatergic transmission within this pathway. Calcitonin gene-related peptide (CGRP) facilitates PB◊CeA excitatory transmission through postsynaptic PKA- and NMDA receptor-dependent mechanisms (Han et al., 2010; Han et al., 2005; Okutsu et al., 2017). Recent work using endogenous peptide release further supports interactions between CGRP and glutamatergic signaling at PB→CeA synapses (Lorsung et al., 2025). Consistent with these findings,chemogenetic inhibition of the PBN→CeA pathway reverses injury-induced pain sensitization whereas chemogenetic activation of the pathway is sufficient to induce pain hypersensitivity in the absence of injury (Torres-Rodriguez et al., 2024). However, these studies do not establish whether VGluT2-mediated glutamatergic transmission specifically from elPBN neurons is required for pain sensitization. Moreover, whether glutamatergic signaling contributes to restraint-stress responses that depend on elPBN PACAP remains unknown.

In the present study, we investigated the contributions of PACAP and glutamatergic signaling from elPBN neurons to pain- and stress-related behaviors. We first tested the contribution of PACAP to nerve injury-induced pain sensitization, and inflammation-induced spontaneous pain-related behaviors using constitutive and elPBN-specific PACAP deletion. In constitutive PACAP knockout mice, we also evaluated CeA-PKCδ neuron recruitment following subthreshold pinch on injured hind paw. We then used optogenetic-assisted circuit mapping to examine glutamatergic transmission from PACAP-expressing elPBN neurons to the CeA and evaluated the consequences of VGluT2 deletion in elPBN neurons across the same pain paradigms. Finally, we asked whether VGluT2-mediated glutamatergic signaling contributes to a restraint stress– induced hypophagia previously shown to depend on elPBN PACAP. By comparing PACAP and glutamatergic signaling across these behavioral contexts, we sought to determine whether these neurochemical signals make overlapping or distinct contributions to the diverse functions of the elPBN.

## METHODS AND MATERIAL

### Animals

Male and female mice (wild-type, transgenic, or knockout) aged 8-17 weeks with a C57BL6 background were housed separated by sex in groups of 2–5 per cage with unlimited access to food and water. Experiments were performed in two different labs with different housing conditions (standard 12-hour light/12-hour dark cycle and reverse 12-hour dark/12-hour light cycle). Housing conditions per experiment are detailed below. For the slice electrophysiology and stress-induced hypophagia experiments, Adcyap1-2A-cre mice (Cre expression restricted to PACAP-expressing neurons of adult mice) and VGluT2^fl/fl^ mice were bred and housed under a standard 12-hour light/12-hour dark cycle (light on from 6 am to 6 pm). For the pain experiments, PACAP^fl/fl^ and VGLuT2^fl/fl^ mice were bred and maintained under the standard light cycle until they underwent stereotaxic surgery. After surgery, these mice were transferred to a reverse 12-hour dark/12-hour light cycle (light on from 9 pm to 9 am) and allowed to entrain to the new cycle for at least 2 weeks prior to any behavioral experiment. PACAP knockout (KO) and wild-type (WT) mice were bred under both lighting conditions. Those originally housed under the standard light cycle were transferred to the reverse light cycle and allowed to entrain to the new cycle for at least 2 weeks prior to experiments. No overt differences in behavioral readouts were noted between mice housed in standard vs reverse light cycle.

All animal care procedures received approval from the National Institute of Mental Health and National Institute of Neurological Disorders and Stroke (NIMH and NINDS) Institutional Animal Care and Use Committees (IACUCs) and adhered to National Institute of Health (NIH) guidelines. The generation and characterization of PACAP-deficient (PACAP^-/-^) mice and conditional PACAP^fl/fl^ strain have been previously described (Bakalar et al., 2022; Hamelink et al., 2002). Pauli et al. (2022) extensively characterized the Adcyap1-2A-Cre line (JAX #030155), demonstrating that its reporter expression produces an anatomical distribution consistent with endogenous *Adcyap1* expression patterns. Heterozygous PACAP^+/-^ male and female mice were crossed to generate PACAP^-/-^ (PACAP-KO) and WT littermates. PACAP genotypes were determined using DNA extracted from tail biopsies. Genotyping was performed by PCR (Transnetyx) using the following primers: CAATTTAGCTAGAGTCATGTATAGTGTAATC (forward primer) and TAAAGCGCATGCTCCAGACTGC (reverse primer). Homozygous PACAP^fl/fl^ mice were used to establish breeding colonies for this strain. Offspring were genotyped using the following primers: Forward, 5′-CCAGAAAACTGGGAGTGGAA-3′; and reverse, 5′-AGACACATTGAGTCGCTGAG-3′. Adcyap1-2A-Cre mice were obtained from The Jackson Laboratory (stock Cat#030155) and were bred with C57BL/6J WT mice to maintain the colony. VGluT2^fl/fl^ mice were sourced from The Jackson Laboratory (STOCK Slc17a6/J, stock Cat#012898). Homozygous VGluT2^fl/fl^ mice were used to establish breeding colonies. Genotyping was performed according to protocols provided by The Jackson Laboratory, and tail samples were submitted to Transnetyx for genotyping.

### Stereotaxic injections

AAV9.hSynap.HI.eGFP-Cre.WPRE.SV40 (Addgene Cat#105540-AAV9) and AAV9.hSynap.HI.eGFP.WPRE.bGH (Addgene Cat#105539-AAV9) were obtained from Addgene (Watertown, MA) and were originally provided as a gift from the James M. Wilson Lab (University of Pennsylvania). Surgical procedures and viral injections were performed in accordance with the NIH-ARAC guidelines for rodent survival surgery. To selectively eliminate PACAP or VGluT2 expression in the external lateral parabrachial nucleus (elPBN), 0.2 µl of AAV9.hSynap.HI.eGFP-Cre.WPRE.SV40 was injected slowly (∼0.1 µl/min) into the elPBN (coordinates: AP −5.2 mm, ML 1.5 mm, DV 3.6 mm) of PACAP^fl/fl^ or VGluT2^fl/fl^ mice. Injections were performed unilaterally in the right hemisphere for pain studies and bilaterally for stress studies. Control mice received 0.2 µl of AAV9.hSynap.HI.eGFP.WPRE.bGH, injected into the elPBN using the same coordinates and unilateral or bilateral injection paradigms. Mice were allowed to recover for 4–6 weeks following viral injection before undergoing restraint stress or sciatic nerve cuff implantation.

For slice electrophysiology experiments, Adcyap1-2A-Cre (PACAP-Cre) mice received a unilateral injection of AAV5-EF1a-DIO-hChR2(H134R)-eYFP (200 nl; ∼1×10¹² infectious particles/ml; UNC Vector Core, Cat#156229; stock titer: 4.0×10¹² GC/ml) into the right elPBN (AP −5.2 mm, ML 1.5 mm, DV 3.6 mm). This viral construct was used to enable selective optogenetic activation of PACAP-expressing elPBN neurons and their projections during electrophysiological recordings.

### Sciatic cuff implantation

The sciatic cuff implantation procedure was conducted on mice 2-4 weeks after the brain surgeries, following the methodology outlined in Benbouzid et al (2008)(Benbouzid et al., 2008). The mice were initially anesthetized with 2% isoflurane at a flow rate of 0.5 L/min. An incision of approximately 1 cm was made in the lateral left thigh’s proximal one-third, exposing and gently stretching the sciatic nerve using forceps. In the group receiving the cuff implant, a 2 mm-long polyethylene tubing (PE20) with a 0.38 mm inner diameter and 1.09 mm outer diameter was carefully placed around the sciatic nerve and then returned to its original position. Following the procedure, the incision on the thigh was closed with wound clips, and the mice were given at least one week to recover before conducting any behavioral experiments.

### Nociceptive Behaviors

Male and female mice were used for study, with behavioral tests on male and female mice conducted on separate days, under red light, during the dark period (9 am to 6 pm). Mice were handled for 3-4 days before behavioral testing using cupping method as described by Hurst and West (2010)(Hurst & West, 2010). Experimenter was blind to treatment conditions during behavioral tests, and each cohort was counterbalanced to include mice from control and experimental groups. The von Frey and acetone test were conducted 6-7 days after cuff implantation on the same day, waiting 30 minutes between tests. The behavioral test boxes were custom made from opaque plexiglass, were ventilated, and measured 11×11×13 cm (NIH Section on Instrumentation). The timeline for the behavioral experiments relative to cuff implantation was as follows: von Frey and Acetone test: 7 or 8^th^ day; Hargreaves test: 8 or 9^th^ day; Randall-Selitto test: 9-12^th^ days. No overt sex differences were seen; thus, data from both sexes were pooled.

#### Acetone test

The acetone test is a commonly used method for assessing sensitivity to cold stimuli (Choi, 1994). Briefly, mice were individually acclimated for 2-3 hours in a ventilated opaque white plexiglass testing chamber placed on an elevated platform with a wire mesh floor. A small drop of acetone was gently applied to the center of the plantar surface of the hind paw. The mice’s nociceptive responses were evaluated for 60 seconds, utilizing a modified 0-2-point system (Colburn et al., 2007). According to this scoring system: a score of 0 represented rapid, transient lifting, licking, or shaking of the hind paw that subsided immediately; a score of 1 indicated lifting, licking, and/or shaking of the hind paw that persisted beyond the initial application but subsided within 5 seconds; a score of 2 represented prolonged, repeated lifting, licking, and/or shaking of the hind paw. The test was repeated 5 times for each hind paw, with a minimum 5-minute interval between tests to allow for recovery. The average of five trials was calculated to determine the threshold value for each hind paw.

#### von Frey filament test

To evaluate mechanical hypersensitivity, paw withdrawal responses to von Frey filaments (North Coast Medical, Inc., San Jose, CA) were measured following the protocol described by Wilson et al. (2019)(Wilson et al., 2019). Mice were acclimated on a mesh floor in individual ventilated opaque white plexiglass chambers for a minimum of 2 hours. Once acclimated, each von Frey filament was applied to the center of the plantar surface of the hind paw for a duration of 2-3 seconds, applying enough force to cause slight bending against the paw. The smallest filament that elicited a paw withdrawal response in at least three out of five measurements was considered as the mechanical threshold for that trial. The average of five trials was calculated to determine the threshold value for each hind paw.

#### Hargreaves test

To assess sensitivity to heat stimulation of the paw, a modified version of the Hargreaves method (Hargreaves et al., 1988; Wilson et al., 2019) was employed. Mice were acclimated in small plexiglass testing chambers with a glass floor maintained at 30°C for 30 min to 1 hour or until they exhibited signs of calmness. Following the habituation period, a noxious radiant light beam, generated by an apparatus (IITC Life Sciences, Woodland Hills, CA), was directed onto the midplantar surface of the hind paw through the glass floor. The latency of paw withdrawal from the radiant light beam was recorded, with an active intensity of 25% and a cutoff time of 15 seconds to prevent tissue damage. The threshold for each hind paw was determined by calculating the average of five trials conducted at 5-minute intervals.

#### Randall-Selitto test

To assess the mechanical pressure threshold of the hind paws, the Randall-Selitto test (Randall, 1957) was performed on lightly anesthetized mice. This test involves gradually applying increasing force to the plantar surface of the hind paw until the animal responds by withdrawing its paw or vocalizing. Initially, mice were anesthetized using 5% isoflurane in an induction chamber. Subsequently, the animals were maintained under light anesthesia with 0.5-1% isoflurane at a flow rate of 0.5 L/min. A maximal cut-off of 200-g force was applied to prevent any tissue damage. Five trials per animal were recorded and the average was calculated and used as threshold for each paw.

#### Nociceptive affective-motivational behavioral assays

To assess affective-motivational responses, we employed a modified version of the protocol outlined by Corder et al. (2019) (Corder, 2019). For mechanical stimulation, we used two von Frey filaments: 0.16 g for light touch and 1.0 g for noxious touch. Each filament was applied to the center of the plantar surface of the hind paw for 2–3 seconds, and behavioral responses were recorded. For heat stimulation, a controlled noxious radiant light beam, was directed onto the midplantar surface of the hind paw through the glass floor. The mice’s aversion scores were evaluated using a 0–3 point system defined as: 0 = No response; 1 = Lifting; 2 = Lifting, shaking, and/or licking; 3 = Escape (jumping or running while lifting). The aversion score for each hind paw was determined by calculating the average of 10 stimulations for mechanical-induced or 5 stimulations for heat-induced nociceptive affective-motivational behavior, respectively.

#### Formalin test

The formalin test is used as a model of inflammatory pain in rodents. In this test, mice received 10 µL of 3% formalin subcutaneously on the plantar surface of either their left or right hind paw. Subsequently, mice were returned to the testing chamber. Time spent performing nociceptive behaviors (including licking, lifting, and shaking of the formalin-injected paw) was recorded in 5-minute bins for 1-hour. For analysis, formalin nociceptive behavior was divided into two phases: Phase 1, 0-10 minutes post-formalin injection, and Phase 2, 10-60 minutes post-formalin injection. To assess paw edema as an indirect measurement of inflammation induced by formalin, we measured dorsoventral diameter of the intraplantar region of the hind paw using a digital caliper (Fisherbrand, Thermo Fisher Scientific).

### Locomotion and Anxiety-like behavior

#### Open field test

The open field test (OFT) was performed in a custom-made square box with dimensions 48×48×43 cm made of black plexiglass (NIH Section on Instrumentation) to measure locomotor and exploratory behavior. Mice were gently placed in a corner of the arena and allowed to freely explore for 10 minutes under 20 lux warm light. The total distance traveled in the entire arena and time spent in the center area, defined as 50% of arena, were measured using CleverSys software (version 3.00, CleverSys Inc, Reston, VA).

#### Elevated zero maze

The elevated zero maze (EZM), purchased from CleverSys, is a circular, elevated track divided into quarters, with 2 closed arms and 2 open arms. The round platform was 53 cm in diameter and 14 cm wide, made of blue plexiglass and elevated 60 cm above the floor. The mice were gently placed onto the center of one closed arm and allowed to freely explore for 6 minutes under 20 lux warm light. The time spent in and entries into the open arm were measured using TopScan tracking software (version 3.00, CleverSys Inc, Reston, VA).

#### Sucrose splash test

The sucrose splash test was performed to evaluate grooming and depressive-like phenotypes, as previously described(Francis-Malave et al., 2023; Kalueff & Tuohimaa, 2004; Yalcin et al., 2005). The test involved spraying 2 ml of a 10% sucrose solution onto the dorsal surface of the mice, after which grooming behavior was assessed based on licking, biting, or scratching the fur to clean off the solution. The time spent grooming was recorded for 10 minutes immediately following the application of the sucrose solution. The coat state of the animals was scored 10-minutes after sucrose application using the following scale: 0 = Completely dry and smooth dorsal coat; 1 = Mostly dry and smooth dorsal coat with a few scruffy patches; 2 = Dry and smooth lower back coat but upper back coat is still scruffy and/or humid; 4 = Scruffy and humid dorsal coat: 8 = wet, soaked dorsal coat.

### Restraint stress and body weight

Restraint stress was performed as previously reported (Jiang & Eiden, 2016), In short, mice were individually housed, transferred to the testing room in their home cages, and given at least one hour to acclimate. Over three days (Days 1–3), they were gently handled to minimize handling-related stress before restraint. Tail vein blood sampling was performed on Day 3 (for NRS: non-restraint stress) and Day 4 immediately after 2-hour restraint (for RS 2-hour). Plasma was separated by centrifugation, stored at -80°C, and corticosterone levels were measured using the DetectX Corticosterone ELISA kit (Arbor Assays, Michigan, USA) per the manufacturer’s protocol. On Day 4, mice were restrained in a DecapiCone for two hours. Body weight was recorded daily throughout the experiment. Acute restraint-induced weight loss was calculated as: [(Day 5 weight – Day 4 pre-restraint weight) / Day 4 pre-restraint weight] × 100. For body weight calculation prior to restraint (NRS), 24-hour weight loss was averaged from Days 1-3.

### RNAscope *in Situ* Hybridization

*In situ* hybridization was conducted on elPBN sections using the RNAScope Multiplex Fluorescent V2 Kit (Cat# 323100) from ACD Bio, following the provided instructions. Fresh-frozen mouse brains were sectioned at 12 µm (coronal or sagittal) on a cryostat and mounted onto Superfrost Plus slides (Fisher Scientific). Sections were fixed in 4% paraformaldehyde for 15 minutes at 4°C, then dehydrated through an ethanol series. After hydrogen peroxide and protease treatments, probe hybridization and signal amplification were conducted as per the manual. Probes used were Mm-Adcyap1 (Cat# 405911) and Mm-Slc17a6-C3 (Cat# 319171-C3), visualized with Opal 520 and Opal 570 fluorophores (Akoya Biosciences). Sections corresponding to approximately bregma -5.0 to -5.3 mm were used for image acquisition and analysis, with 1-3 sections quantified and averaged per animal. Images were acquired using a Zeiss Axio Scan (Carl Zeiss Microscope, Thornwood, NY) or Leica Stellaris confocal microscope (Leica Microsystems Inc., Deerfield, IL) at the NIMH Light Imaging Facility. Confocal images were acquired as z-stacks with a typical step size of 1-2 µm. DAPI staining was used as a nuclear counterstain and was provided in the mounting medium (SlowFade Gold Antifade Reagent with DAPI, Invitrogen, Cat# S36938).

### Immunohistochemistry

#### Injection verification in elPBN and Stress-induced c-Fos

Immunohistochemistry followed methods described by Jiang et al. (2023). In brief, mice were perfused with cold 4% paraformaldehyde in PBS. Brains were fixed in 4% PFA at 4-8°C for 24-hours, then immersed in 30% sucrose/PBS at 4–8°C for 48–72 hours before sectioning. Brains were sliced into 40µm sections using a microtome. Free-floating sections were rinsed in PBS with 0.5% Triton X-100 (PBST; 3 × 15 min), blocked in 10% normal goat or donkey serum in PBST for 1-hour at room temperature, and incubated overnight at 4°C with primary antibodies diluted in blocking solution. The next day, sections were washed in PBST (3×15 min) and incubated with Alexa 555-, Alexa 488-, or Alexa 647-conjugated secondary antibodies (1:300; Millipore Sigma) for 2 hours at room temperature. Sections were mounted using ProLong Gold Antifade Mountant (ThermoFisher Scientific, Waltham, MA). Primary antibodies included rabbit anti-GFP (Invitrogen, Cat# A-11122), rabbit anti-cFos (EnCor Biotechnology Inc., Gainesville, FL, Cat# RPCA-c-FOS), and guinea pig anti-VGluT2 (Synaptic Systems, Cat# 135404). Confocal imaging was performed on a Leica Stellaris microscope (Leica Microsystems Inc., Deerfield, IL) at the NIMH Light Imaging Facility. c-Fos-positive neurons in the extended amygdala were quantified by counting immunoreactive (IR) signals.

#### Pinch induced c-Fos

Mice received either subthreshold (25 g; insufficient to evoke a pain response) or noxious (180-200 g; strong enough to produce observable pain behavior) pinch stimulation using the Randall-Selitto method. Stimuli were delivered at 1 stimulation/minute for 30 minutes. Subthreshold stimulation targeted only the injured hind paw, while noxious stimulation was applied to both hind paws. One hour after pinch stimulation, mice were anesthetized via i.p. injection of 1.25% Avertin (2,2,2-tribromoethanol and tert-amyl alcohol in 0.9% NaCl; 0.025 ml/g body weight) for transcardial perfusion. First, 0.9% NaCl (37°C) was perfused for a few minutes until the blood ran clear. Then, 100 mL of 4% paraformaldehyde in phosphate buffer (PFA/PB), chilled on ice, was perfused at a rate of 20 mL/min. After successful perfusion, brains were removed and post-fixed in 4% PFA/PB overnight at 4°C, followed by cryoprotection in 30% sucrose/PB for 48 hrs at 4°C. Brains were then sectioned using a sliding microtome, secured with a base layer of O.C.T compound (Fisher HealthCare). Coronal slices (30-35 μm) containing the central amygdala and parabrachial nucleus were collected and stored in 1X PBS containing 0.01% sodium azide at 4°C until immunostaining. On the day of the immunostaining, slices were rinsed in 1X PBS and incubated in 0.1% Triton X-100 in PBS for 10-minutes at room temperature. Slices were then blocked in 5% normal goat serum (NGS; Vector Labs, Burlingame, CA) containing 0.1% Triton X-100, 0.05% Tween-20 and 1% bovine serum albumin (BSA) for 30 minutes. Primary and secondary antibodies were diluted in a 1.5% NGS blocking solution containing 0.1% Triton X-100, 0.05% Tween 20 and 1% BSA. Secondary antibody incubation was conducted in the dark to protect from light exposure. Primary antibodies: mouse anti-PKCδ (BD Biosciences, Cat# 610397, 1:1000), rabbit anti-Phospho-Fos (Cell Signaling, Cat# 5348, 1:1000) were incubated for 24 hours at 4°C, followed by 1 hour at room temperature on a shaker. Sections were then rinsed 3 times in 1x PBS and incubated with secondary antibodies: Alexa Fluor 488-goat anti-mouse (Thermo Fischer, Cat# A-11001, 1:500) and Alexa Fluor 647-conjugated goat anti-rabbit (Invitrogen, Cat# A21244, 1:500) for 2 hours at room temperature on a shaker. Slices were mounted on VWR® Superfrost Plus Slides and air-dried overnight in a covered box before applying DAPI Fluoromount-G mounting media (Southern Biotech) and coverslip.

#### Image acquisition and analysis for pinch induced c-Fos

All images were acquired using a Nikon Eclipse Ti-Si laser scanning confocal microscope. An And or Zyla camera was used for all 10x, 20x, and 40x oil-immersion high-magnification images. Laser conditions were kept constant throughout experiments. For low-magnification (2x) images, a Hamamatsu Flash was used. Quantification of c-Fos and PKCδ was performed on z-stacks at 10 x magnification, 5.6 µm step intervals. All images were analyzed using NIS Elements AR software. Images were represented as maximum intensity projections and processed using binary bright spot detection to identify c-Fos- and PKCδ-positive neurons. Each detection was visually verified and manually corrected if necessary. Regions of interest (ROIs) were manually delineated using the Paxinos & Franklin (2008) mouse brain atlas as a reference. Colocalization of c-Fos and PKCδ-positive neurons was assessed using automated binary overlap detection followed by visual confirmation. For quantitative analysis, 10 coronal brain sections per mouse spanning bregma -0.71 mm to -1.79 mm were evaluated. The percentage were calculated by dividing the number of co-labeled (c-Fos+PKCδ) cells by the total number of either the PKCδ+ or c-Fos+ cell populations and multiplying by 100 to determine the proportion of PKCδ+ cells expressing c-Fos and of c-Fos+ cells expressing PKCδ, respectively.

### Slice electrophysiology

#### Acute Slice preparation

Mice were deeply anesthetized with ketamine/xylazine (100 mg/kg ketamine and 10 mg/kg xylazine; *Guide for the Care and Use of Laboratory Animals,* NIH Division of Veterinarian Resources, DVR) and then decapitated. Brains were rapidly removed and placed in ice-cold NMDG-based cutting solution (Ting et al., 2014) containing (in mM): 92 NMDG, 20 HEPES, 25 glucose, 30 NaHCO_3_, 2.5 KCl, 1.2 NaPO_4_ saturated with 95% O_2_/5% CO_2_, with an osmolarity of 303-306 mOsm/L (VAPRO® 5600, *Wescor*). The brain was rapidly block-cut, dried on filter paper, and glued to a platform containing ice-cold NMDG-based cutting solution within a vibratome (VT1200, Leica™). Coronal slices with 250 μm of thickness, containing the CeA were cut at a speed of 0.07 mm/s. Following slicing, sections were incubated in a chamber containing NMDG-based cutting solution for 10-15 min at 34°C. Slices were subsequently transferred to a chamber filled with modified holding aCSF saturated with 95% O_2_/5% CO_2_ containing (in mM): 92 NaCl, 20 HEPES, 25 glucose, 30 NaHCO_3_, 2.5 KCl, 1.2 NaPO_4_ (303-306 mOsm/L) at room temperature for at least 1 hour before starting the recordings.

#### Optogenetic Excitatory Postsynaptic Currents (oEPSCs) recordings

Slices remaining in the holding aCSF solution were transferred to the recording chamber and cells were visualized using infrared differential interference contrast (IR-DIC) optics on an upright microscope (Olympus BX5iWI). The recording chamber was perfused by a peristaltic pump (*World Precision Instruments*) at a flow rate of 1.5-2.0 ml per minute with aCSF solution containing (in mM): 126 NaCl, 2.5 KCl, 1.4 NaH_2_PO_4_, 1.2 MgCl_2_, 2.4 CaCl_2_, 25 NaHCO_3_, and 11 glucose (303-305 mOsm/L) and warmed to 32-34°C by passing it through a feedback-controlled in-line heater (TC-324B; *Warner Instruments*). For whole-cell patch-clamp of oEPSC recordings, we utilized glass microelectrodes (3-5 MΩ) filled with a solution containing (in mM): 117 cesium methanesulfonate, 20 HEPES, 0.4 EGTA, 2.8 NaCl, 5 TEA-Cl, 4 Mg-ATP, 0.4 Na-GTP, and 5 QX-314 (280-285 mOsm/L) and 0.1% of Biocytin (Sigma-Aldrich, Cat# B4261) for identification of recorded neurons. Monosynaptic oEPSCs evoked in CeA-PKCδ neurons (confirmed after the recordings) by stimulation of PACAPergic elPBN ChR2-expressing terminals were isolated with 1 μM tetrodotoxin (TTX, Cayman Chemical, Cat# 4368-28-9) and 4-aminopyridine (4-AP; 50 μM, Cayman Chemical, Cat# 504-24-5). Patch-clamped CeA neurons were monitored in voltage-clamp mode at a membrane potential of −55 mV. If a current of more than 10 pA was detected in response to a brief pulse of blue light (473 nm) by controllable Illumination system (pE-300^ultra^, CoolLED©), the laser power (5-10 mW) or duration (max. 1 ms) was increased until a 50–1200 pA oEPSC was obtained. After the baseline stabilized (>5 min), post-synaptic currents in response to optical stimulation of elPBN terminals were recorded before and 5 min after bath exchange to ACSF containing 10 μM 6-cyano-7-nitroquinoxaline-2,3-dione (CNQX; Cayman Chemical, CAS 115066-14-3). A 8.62 mM stock solution of CNQX was made in water, stored at –20°C and dissolved in ACSF on the day of the experiment. Neurons were recorded utilizing a Multiclamp 700B amplifier (*Molecular Devices*™). Data were filtered at 2 kHz and digitized at 20 kHz using a 1440A Digidata Digitizer (*Molecular Devices*™). Series-resistance (10-20 MΩ) was monitored using a -5 mV voltage step. Cells with >20% change in series resistance were discarded from further analysis. Following electrophysiological recording, the pipette was slowly retracted from the cell, and the slice was transferred to 4% paraformaldehyde overnight and then underwent immunohistochemistry staining.

#### Biocytin Labeling and Immunohistochemistry

Following electrophysiological recordings, slices containing biocytin-filled cells were collected and stored overnight in 4% paraformaldehyde (PFA) for fixation. The fixed slices were washed 4–5 times in Tris-buffered saline containing 0.1% Tween-20 (TBST; approximately 30 min per wash) at room temperature (RT) with gentle shaking and then incubated overnight at 4°C in 10% normal goat serum (NGS) in TBST for blocking. Slices were subsequently incubated with primary antibody diluted in 3% NGS in TBST for 48– 72 h at 4°C with gentle shaking. For PKCδ immunolabeling, purified mouse anti-PKCδ antibody (BD Biosciences, Cat# 610397) was used at a dilution of 1:500. Following primary antibody incubation, slices were washed five times in TBST (approximately 1 h per wash) at RT and incubated with Alexa Fluor 488 goat anti-mouse secondary antibody (1:1000) diluted in 3% NGS in TBST for 48–72 h at 4°C with gentle shaking. Slices were then washed 4–5 times in TBST (approximately 1 h per wash) at RT. For mounting, slices were transferred individually to slides, allowed to air-dry briefly, and coverslipped using VECTASHIELD mounting medium containing DAPI for nuclear counterstaining. Coverslip edges were sealed with nail polish, and slides were stored at 4°C until imaging.

#### Analysis

The oEPSCs traces were processed and analyzed with the offline software Clampfit 11.2 (Molecular Devices™) and the Prism 8.0 (GraphPad), respectively. After the baseline stabilized (>5 min), we conducted experiments to investigate CNQX effects. A train of light pulses at 40 Hz was delivered to obtain a current response (oEPSCs). For each light pulse, we recorded 5 sweeps at 15-second intervals. The sweeps were obtained under aCSF and +CNQX [10 µM] conditions. We then averaged the sweeps and performed threshold analysis on the averaged trace to obtain the maximum amplitude (pA) of each response. We performed a paired t-test to compare the change in maximum amplitude (pA) in the 1st peak between aCSF and +CNQX conditions (p < 0.05).

## Statistical Analysis

All summarized data are expressed as mean± SEM. Statistical analyses were conducted using GraphPad Prism (v. 10.0) with two-way analysis of variance (ANOVA) followed by Sidak’s or Dunnett’s multiple comparison tests, unpaired or paired two-tailed t-test. The n (number of animals), statistical test, and p values are presented in the figure legends and statistical table. The value of p < 0.05 was considered statistically significant.

## Results

### Constitutive knockout of PACAP prevents nerve injury-induced pain sensitization without affecting baseline nociception

To assess the role of PACAP in pain processing, we utilized a constitutive PACAP knockout (KO) alongside wild-type (WT) control mice. Neuropathic pain was induced using the cuff model (Benbouzid et al., 2008), and nociceptive responses were evaluated through a battery of behavioral assays, including acetone (cold), Hargreaves (heat), von Frey (tactile), and Randall-Selitto (pressure) tests (**Figure 1A**). In WT mice, sciatic nerve cuff implantation resulted in pain-related hypersensitivity in the injured hind paw across all tested modalities (cold, tactile, pressure, and heat) consistent with previous findings (Wilson et al., 2019). However, PACAP-KO mice displayed little (tactile) to no signs (all other tests) of cuff-induced hypersensitivity (**Figure 1B**). Compared to WT controls, PACAP-KO mice exhibited significantly reduced responsiveness to acetone (p < 0.0001), higher withdrawal thresholds to tactile stimuli (p < 0.0001), increased tolerance to pressure (p < 0.0001), and prolonged withdrawal latencies to heat stimulation (p < 0.0001) in the paw ipsilateral to cuff implantation. Notably, evoked responses to cold, heat, and pressure in the paw ipsilateral to cuff implantation of PACAP-KO mice were comparable to those of the contralateral paw, indicating that knocking out PACAP completely blocked nerve injury-induced hypersensitivity to these modalities. There was a small but significant (p < 0.01) difference in withdrawal thresholds to tactile stimuli, suggesting that there is a small residual hypersensitivity to tactile in the KO mice. Behavioral responses in the uninjured side were unaffected in PACAP-KO mice, compared to WT across all modalities, demonstrating that knocking out PACAP does not affect baseline nociception (**Figure 1B**).

**Figure 1:**
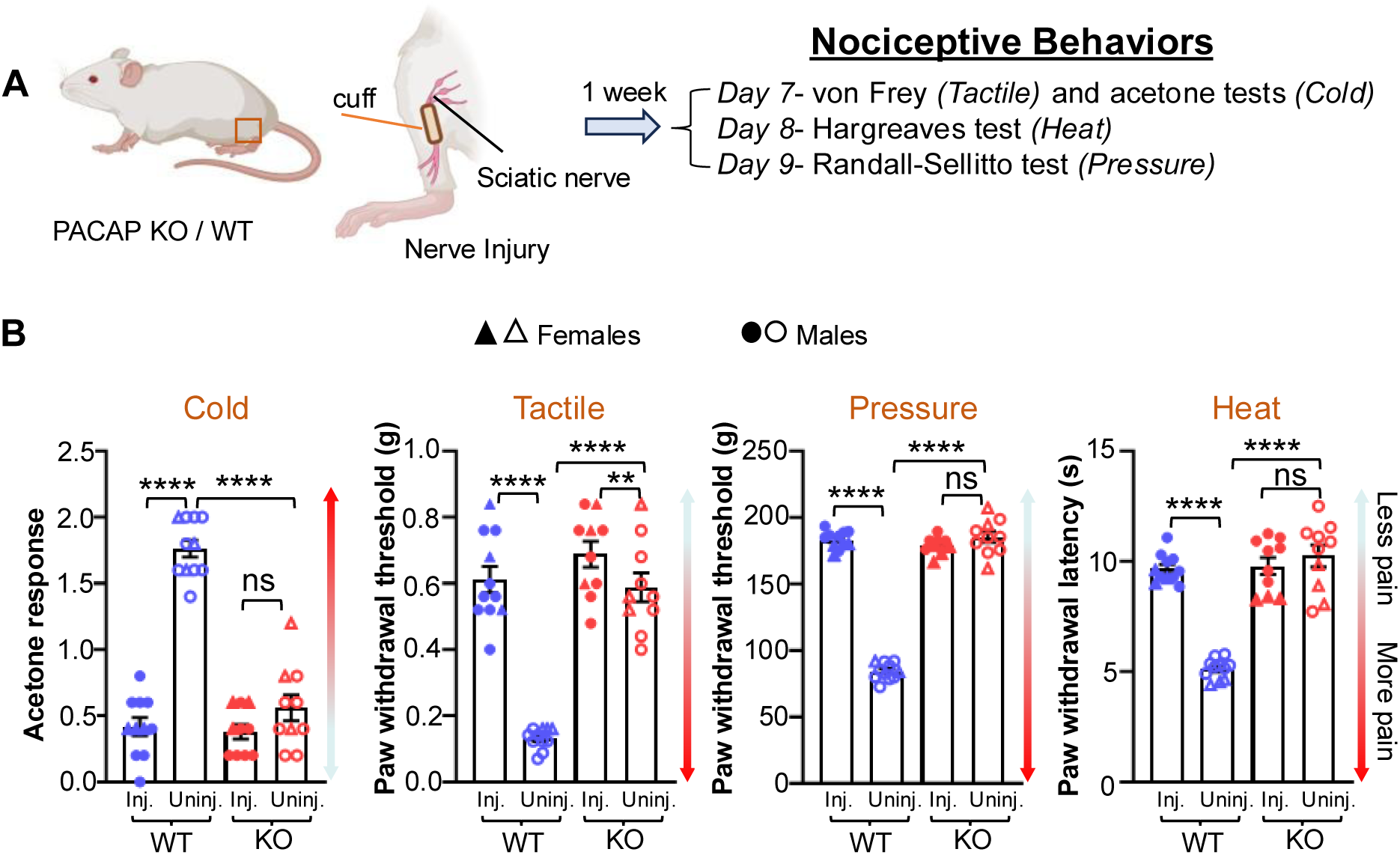
Constitutive knockout of PACAP prevents injury-induced pain sensitization. (A) Experimental design: One-week post-cuff implantation in the sciatic nerve, a series of behavioral assays were conducted to measure evoked nociceptive responses in constitutive PACAP knockout (KO) and wildtype (WT) control mice. (B) Nociceptive responses in the hind paws of WT and constitutive PACAP-KO mice across different modalities—cold, tactile, heat, and pressure. Two-way repeated measure ANOVA followed by Sidak’s post-hoc test; **p < 0.01; ****p < 0.0001. Sample size: n = 11 WT mice (8 males, 3 females) and 10 constitutive PACAP-KO mice (7 male, 3 females). WT= wild-type, KO= knockout, ns= non-significant, Inj.= injured paw, Uninj. = uninjured paw. Individual mice are represented by scatter points. Triangle symbols indicate female and circle indicate male mice. All data are presented as mean ± SEM.

### PACAP deletion restricted to elPBN neurons phenocopies the pain-related effects seen in constitutive PACAP knockout mice

elPBN neurons are crucial for processing and transmitting pain in injured states (Allen et al., 2023; Bernard, 1990; Chiang et al., 2020; Torres-Rodriguez et al., 2024) and PACAP in these neurons are required for some stress-related behaviors (Jiang et al., 2023; Kang et al., 2024). To investigate the contribution of PACAP in elPBN neurons in pain processing, we generated animals with PACAP deletion anatomically restricted to elPBN, as described by Jiang et al. (2023). To achieve localized PACAP deletion in elPBN neurons, we stereotaxically injected Cre-expressing virus (AAV9.hSyna-eGFP-Cre) or a control GFP virus (AAV9.hSyn-eGFP) into the elPBN of PACAP^fl/fl^ mice (**Figure 2A**). Both cre-expressing and GFP-expressing virus injections were localized within the elPBN (**Figure 2A**). To confirm PACAP deletion in the elPBN, we performed in-situ hybridization (ISH) for *Adcyap1* (encoding PACAP). These experiments revealed markedly attenuated PACAP mRNA expression in elPBN neurons from mice infected with Cre-expressing virus when compared with the contralateral, uninfected elPBN within the same animals or with control mice injected with GFP-expressing virus (**Figure 2B-C**). Deletion of PACAP did not significantly alter the mRNA expression of *Slc17a6* (encoding VGluT2), demonstrating that our experimental strategy selectively deletes PACAP in the elPBN.

**Figure 2:**
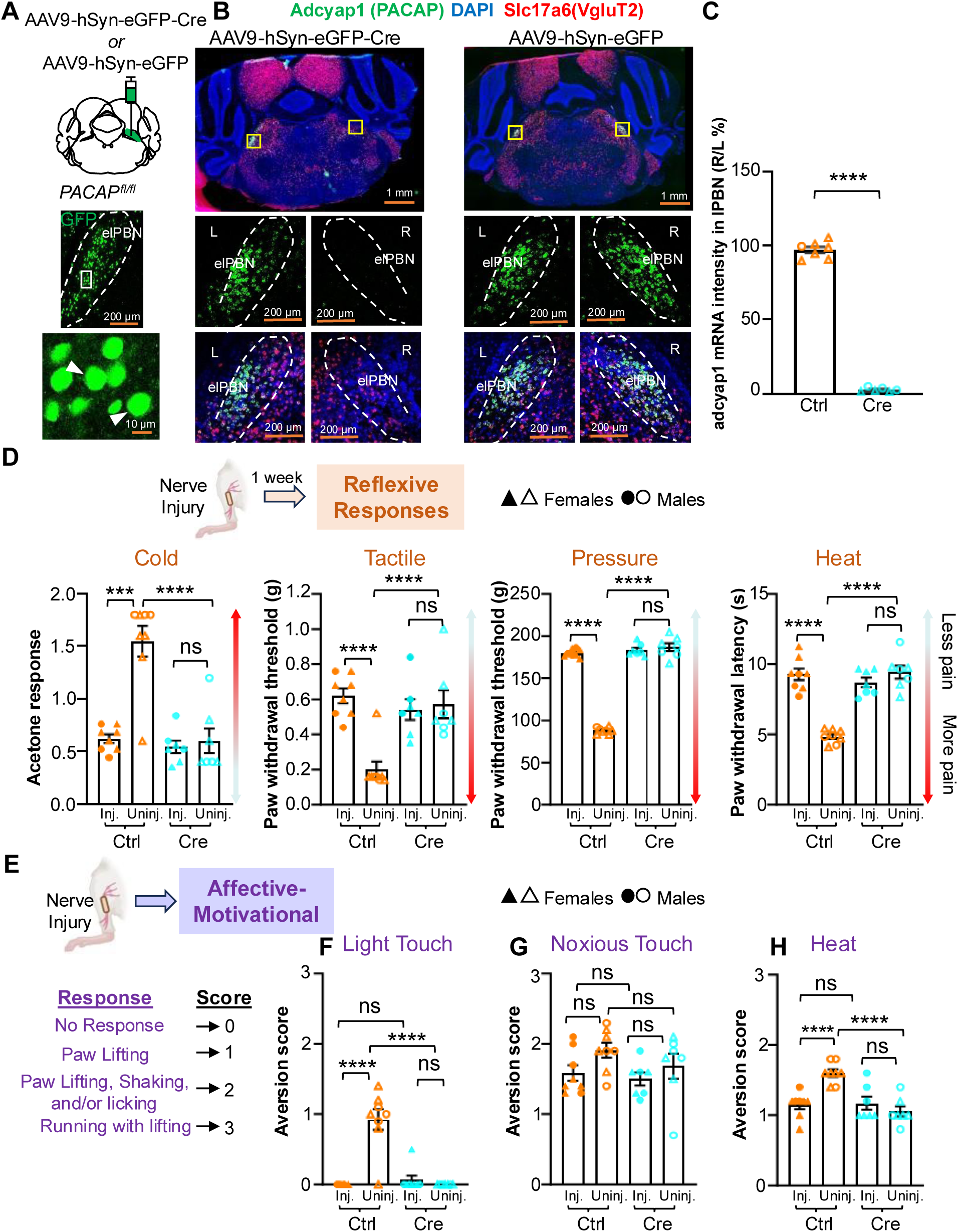
PACAP deletion in elPBN neurons reduces injury-induced pain sensitization and affective-motivational responses. (A) Experimental design: deletion of PACAP in elPBN neurons in PACAP^fl/fl^ mice was achieved by injecting AAV9-hSyn-eGFP-Cre or corresponding control AAV9-hSyn-eGFP virus into the right elPBN. Representative images of coronal slices from mouse brain injected with AAV9-hSyn-eGFP-Cre (middle panel); higher magnification image shows transduced cells (lower panel). Scale bars represent 200 µm for low magnification and 10 µm for high magnification images. Solid arrowhead indicates transduced cells. (B) Evidence of PACAP deletion in elPBN neurons. Top panel: representative coronal brain slice images from an in-situ hybridization experiment targeting the parabrachial area. PACAP^fl/fl^ mice were injected with either Cre-expressing virus (left panel) or corresponding control virus (right panel) into the right elPBN neurons. PACAP mRNA (Adcyap1) is shown green, and VGluT2 mRNA (Slc17a6) is shown in red. Yellow boxes highlight the elPBN region on both the left and right sides of brain. Middle panel showing magnified images of the yellow boxed regions from top panel, indicating PACAP mRNA expression, labeled as left (L) and right (R) in elPBN neurons. Lower panel depicting co-labeled signals of PACAP and VGluT2 mRNA in elPBN neurons on both the left and right sides. Scale bars represent 1 mm for low magnification and 200 µm for high magnification images. (C) The Adcyap1 mRNA signals in the elPBN were quantified and the results were expressed as a percentage ratio of the right side to the left side (R/L %). Sample sizes: PACAP RNAscope, n=7 control mice (1 male, 6 females) and 6 (3 males, 3 females) Cre viral injected mice. ****p < 0.0001, student’s t-test. (D-E) Injury-induced behavioral response: (D) Reflexive responses for different sensory modalities, and (E) affective-motivational responses to light touch (F), noxious touch (G) and noxious heat (H) in PACAP^fl/fl^ mice injected with either AAV9-hSyn-eGFP-Cre or AAV9-hSyn-eGFP virus in the right elPBN. Two-way repeated measure ANOVA followed by Sidak’s post-hoc test; ***p < 0.001; ****p < 0.0001. n = 8 control mice (3 males, 5 females) and 7 Cre mice (3 males, 4 females). Ctrl= control virus AAV9-hSyn-eGFP, Cre= AAV9-hSyn-eGFP-Cre virus, ns= non-significant, Inj.= injured paw, Uninj. = uninjured paw. Scatter points represent individual mice. Triangle symbols indicate female and circle indicate male mice. All data are presented as mean ± SEM.

After a week of sciatic nerve cuff implantation, we conducted a battery of nociceptive behavioral tests to evaluate injury-induced hypersensitivity (**Figure 2D**). As seen in the constitutive PACAP-KO mice, Cre-mediated PACAP deletion in the elPBN significantly reduced cuff-induced cold (p < 0.0001), tactile (p < 0.0001), pressure (p < 0.0001), and heat (p < 0.0001), hypersensitivity in the hind paw ipsilateral to nerve injury compared to control mice (**Figure 2D**). Behavioral responses in the injured paw of elPBN PACAP deletion mice were comparable to those in the uninjured contralateral paw, demonstrating that PACAP deletion in the elPBN completely blocks hypersensitivity after cuff implantation in the sciatic nerve. Responses in the hind paw contralateral to nerve injury were unaffected by PACAP deletion in the elPBN, confirming that PACAP does not contribute to baseline nociception (**Figure 2D**).

In addition to the reflexive paw withdrawal, noxious stimuli elicit protective and escape-seeking behaviors such as guarding, running, and jumping (Corder, 2019). These non-reflexive actions have been proposed to represent the affective-motivational component of pain and rely on brain circuits that can be distinct from those mediating reflexive responses (Huang et al., 2019). To assess the contribution of PACAP in the elPBN to affective-motivational response to hind paw stimulation, we monitored protective and escape-seeking behaviors during paw stimulation in control mice and mice with PACAP deletion in the elPBN (**Figure 2E**). In control mice, aversion scores in response to light touch or heat were significantly (p < 0.0001) higher in the paw ipsilateral to nerve injury than the contralateral paw (**Figure 2F and 2H**). Responses to noxious touch, however, were comparable in both hind paws (**Figure 2G**). Mice with PACAP deletion in the elPBN exhibited significantly (p < 0.0001) lower aversion scores in response to light touch and heat stimulation of the paw ipsilateral to nerve injury compared to control mice (**Figure 2F and 2H**), demonstrating that PACAP in the elPBN contributes to the affective-motivational component of pain-related responses to these modalities. Aversion scores in response to light touch and heat stimulation of the paw ipsilateral to nerve injury were comparable to those seen after stimulation of the contralateral hind paw, demonstrating a complete blockade of these affective-motivational responses by PACAP deletion in the elPBN. Notably, responses to noxious touch were comparable in all mice regardless of treatment (**Figure 2G**), consistent with the lack of PACAP contribution to baseline nociception. Furthermore, no significant difference in aversion scores was observed in the uninjured paw contralateral to nerve injury in mice with PACAP deletion in the elPBN when compared to control mice in any of the modalities tested (**Figure 2F-H**). Together, these findings demonstrate that PACAP deletion restricted to elPBN neurons phenocopies the blockade of reflexive pain-related responses observed in constitutive PACAP-KO mice. These results further indicate that PACAP in the elPBN neurons contributes to affective-motivational responses associated with nerve injury-induced hypersensitivity, but not to affective-motivational responses in uninjured states.

### Constitutive knockout of PACAP reduces recruitment of CeA-PKCδ neurons following subthreshold pinch of the hind paw ipsilateral to nerve injury

The elPBN is the major source of PACAPergic inputs to the CeA (Boucher et al., 2022; Jiang et al., 2023; Missig et al., 2014). Constitutive PACAP knockout or PACAP deletion in the elPBN abolishes restraint stress-induced expression of the neuronal activity marker c-Fos in CeA neurons that express protein kinase C delta (CeA-PKCδ) (Jiang et al., 2023). c-Fos expression after noxious stimulation (pinch) of the hind paw has also been shown to be localized to CeA-PKCδ neurons (Wilson et al., 2019). We evaluated whether pinch-induced c-Fos in CeA-PKCδ neurons is also dependent on PACAP using constitutive PACAP-KO mice and c-Fos immunofluorescence staining (**Figure 3A**). Based on our behavioral experiments showing that PACAP contributes to nerve injury-induced hypersensitivity but not baseline responses to noxious stimulation (**Figures 1-2**), we predicted that PACAP modulation of c-Fos expression in CeA-PKCδ neurons would also be specific to hypersensitive states. To test this hypothesis, we applied subthreshold pinch stimulation to the hind paw ipsilateral to nerve injury in PACAP-KO mice and WT littermate controls and evaluated c-Fos expression in the CeA (**Figure 3A**). After subthreshold pinch stimulation (**Figure 3B**), the total numbers of PKCδ-positive and c-Fos-positive cells in the CeA were comparable between PACAP-KO and WT mice (**Figure 3C-D**). However, colocalization analysis revealed significantly (p < 0.01) fewer neurons co-expressing c-Fos and PKCδ in PACAP-KO mice compared to WT mice across the rostro-caudal extent of the CeA (**Figure 3E**). When normalized, the proportion of c-Fos-positive cells that express PKCδ was comparable between genotypes (**Figure 3F, left).** In contrast, the proportion of PKCδ neurons expressing c-Fos was significantly (p < 0.01) lower in PACAP-KO mice compared with WT mice (**Figure 3F, right)**, indicating reduced recruitment of CeA-PKCδ neurons following subthreshold stimulation in the absence of PACAP. Following noxious pinch stimulation (**Figure 3G**), the total numbers of PKCδ-positive cells, c-Fos-positive cells, and co-labeled neurons were comparable between WT and PACAP-KO mice across the rostro-caudal extent of the CeA (**Figure 3H-J**). Likewise, normalized measures of c-Fos and PKCδ colocalization were also comparable between genotypes (**Figure 3K**). Together, these findings indicate that PACAP contributes to the recruitment of CeA-PKCδ neurons following subthreshold stimulation in the nerve-injured state, whereas c-Fos recruitment following strongly noxious stimulation was not detectably altered by constitutive PACAP loss.

**Figure 3:**
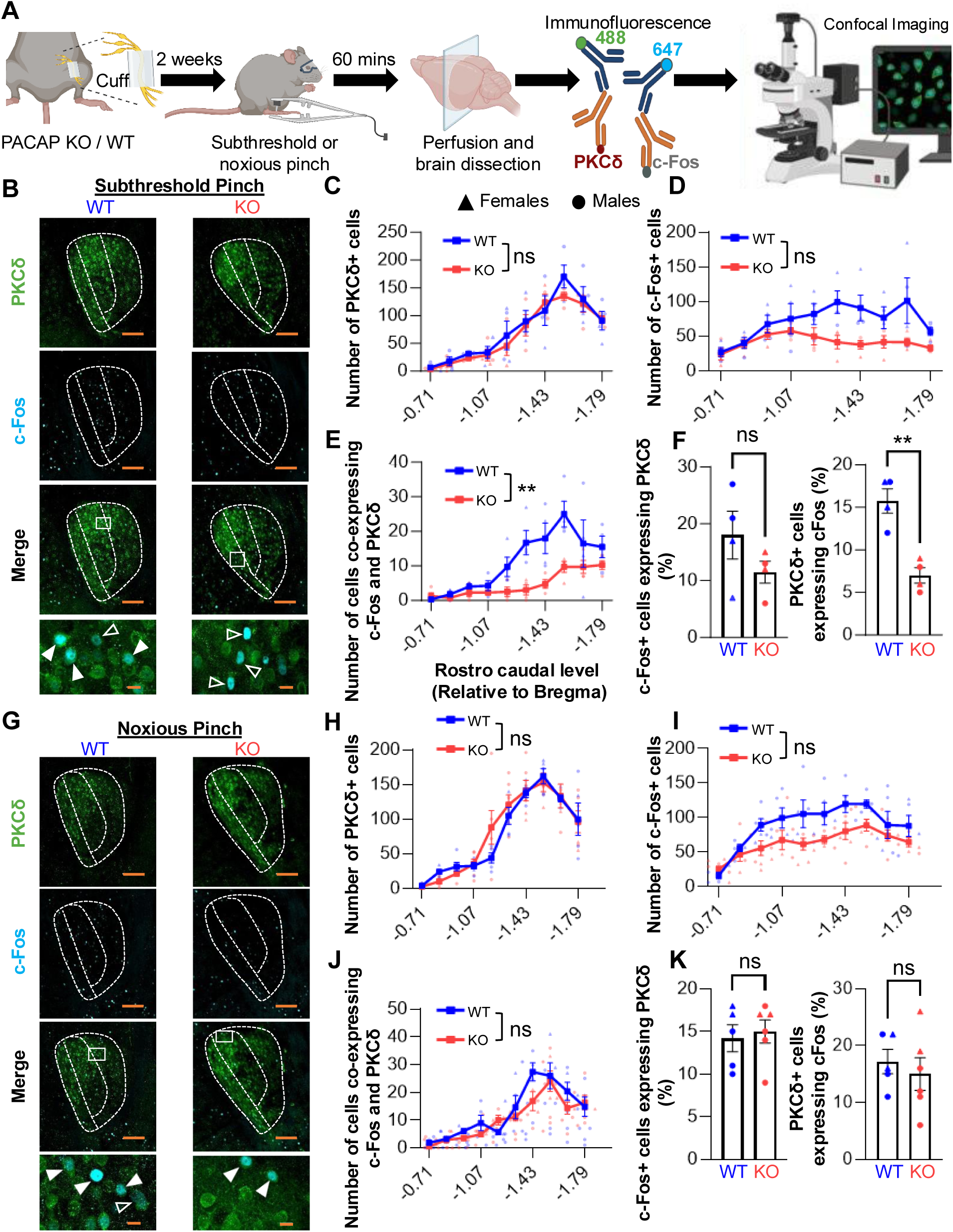
Constitutive knockout of PACAP decreases pain sensitization-induced c-Fos expression in CeA-PKCδ neurons. (A) PACAP-KO and WT mice underwent cuff implantation on the left hind paw. Two weeks later, mice received a cFos induction procedure: either a subthreshold 25 g pinch to the cuffed hindpaw (subthreshold stimulus group) or a noxious (180-200g) bilateral pinch (noxious pinch group), followed by transcardial perfusion for histological analysis. Coronal brain sections were processed for immunohistochemistry, with c-Fos and PKCδ labeling. (B-K) Representative images of the right CeA from subthreshold pinch (B) or noxious pinch (G) animals immunostained for PKCδ (green), c-Fos (cyan), and their colocalization (merge). Lower panels depict magnified images with solid arrowheads indicating cells co-expressing PKCδ and c-Fos, and open arrowhead indicating single labeled c-Fos+ cells. Quantification of PKCδ+ cells (C and H), c-Fos+ cells (D and I), and cells co-expressing PKCδ and c-Fos (E and J) through the rostro-caudal CeA. (F and K) Mean ± SEM percentage of c-Fos+ cells co-labeled with PKCδ (left), and PKCδ+ cells co-labeled with c-Fos+ (right). n = 4 control mice (males 2, females 2) and 4 PACAP-KO mice (males 2, females 2) for subthreshold pinch and 5 control mice (males 3, females 2) and 6 PACAP-KO mice (males 4, females 2) for noxious pinch. Unpaired two-tailed t-test; **p < 0.01; ns = non-significant. All data are presented as mean ± SEM. Scale bars represent 100 µm on lower magnification images and 10 µm for higher magnification images.

### Optogenetic activation of PACAPergic elPBN terminals elicits glutamatergic excitatory postsynaptic responses in CeA-PKCδ neurons

PACAPergic neurons of elPBN are also glutamatergic (Zhang et al., 2021) and show dense terminal labeling in the CeA, selectively surrounding the cell bodies of CeA-PKCδ neurons (Jiang et al., 2023). Several studies have further shown that glutamatergic transmission in the elPBN◊CeA pathway is potentiated in the context of persistent pain (Neugebauer, 2002). To determine whether glutamate is also released by PACAPergic terminals in the CeA, we used optogenetically assisted circuit mapping in brain slices containing the CeA of PACAP-Cre mice stereotaxically injected into the elPBN with an AAV virus (AAV5-EF1a-DIO-hChR2(H134R)-eYFP) expressing channelrhodopsin (ChR2) only in cells expressing Cre (**Figure 4A**). Whole-cell voltage-clamp recordings from CeA neurons were conducted 6 weeks after the virus injection in the elPBN (**Figure 4B**). Excitatory postsynaptic currents (EPSCs) in response to light stimulation of PACAPergic elPBN terminals were observed in 7 CeA neurons (**Figure 4C**). Bath application of the glutamate receptor antagonist CNQX (6-cyano-7-nitroquinoxaline-2,3-dione; 10 µM) significantly (p < 0.05) reduced the amplitude of light-evoked EPSCs (n=6), demonstrating that PACAPergic elPBN terminals also release glutamate in the CeA (**Figure 4C-D**). At the end of recordings, neurons were filled with biocytin for subsequent identification. Immunostaining for PKCδ on the recovered cells showed that 6/7 cells were PKCδ+. These combined results demonstrate that PACAPergic elPBN terminals also release glutamate and can elicit postsynaptic responses in CeA-PKCδ neurons.

**Figure 4:**
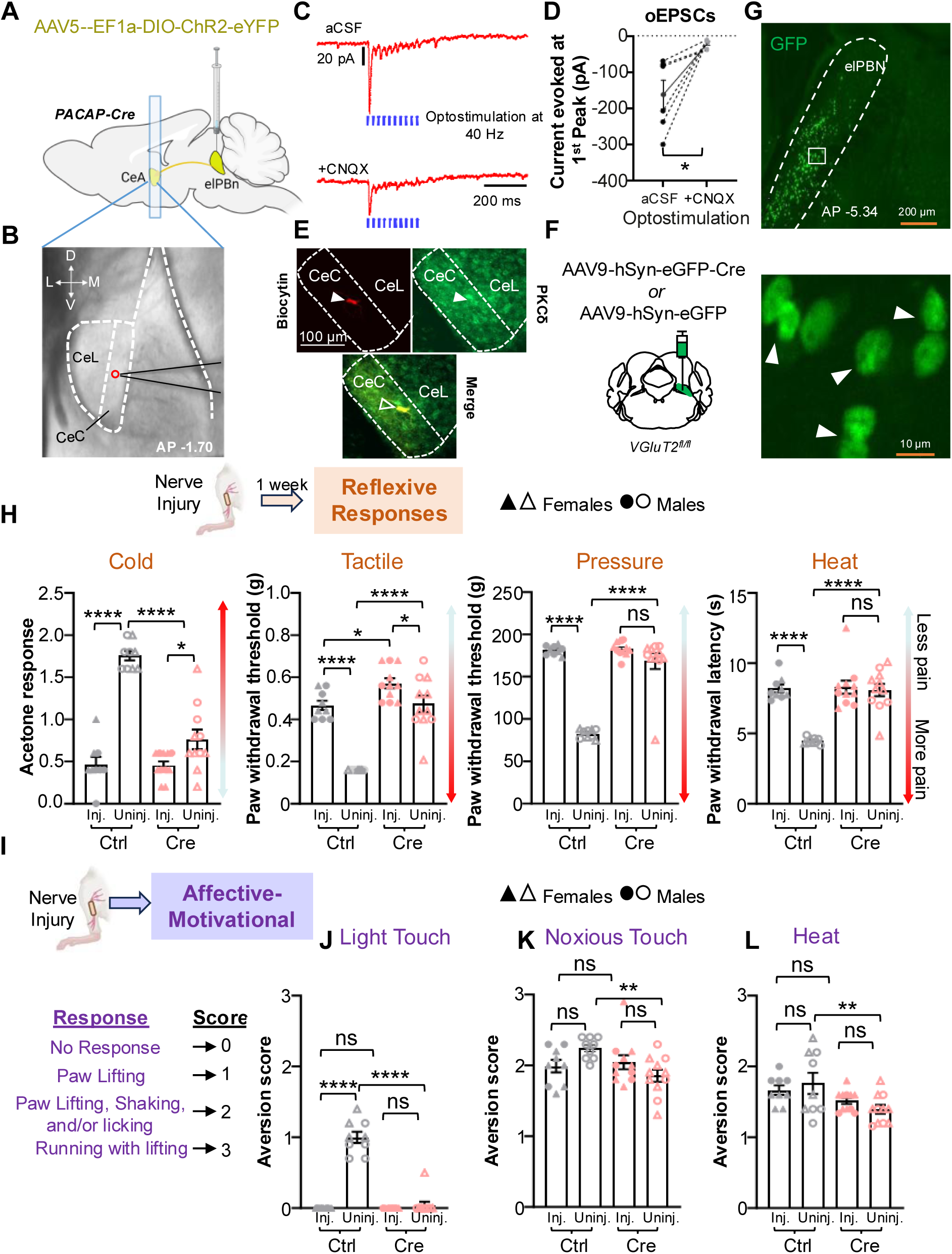
elPBN PACAPergic inputs to CeC PKCδ⁺ neurons are CNQX-sensitive and contribute to injury-induced pain sensitization and affective-motivational responses. (A-E) **Excitatory post-synaptic responses to optical stimulation of** elPBn^PACAP^ terminals in the CeA are CNQX-sensitive. (A) AAV5-EF1a-hChR2(H134R)-eYFP was injected into elPBn of PACAP-Cre mice, and recordings were obtained after 6 weeks. (B) Recording site of PKCδ positive cell in the CeA. Bregma AP: -1.70. (C) Representative traces of optically evoked postsynaptic currents (oEPSCs) in a CeA-PKCδ cell. (D) Quantification showing the amplitude of the oEPSC decreased with [10µM] of CNQX application into the bath. (n = 6 cells from 2 mice; 1 male and 1 female). (E) Representative images of a coronal slice depicting a biocytin-filled CeA neuron (upper left; red) recovered and stained for PKCδ (upper right; green) after recording, confirming that the recorded neuron was PKCδ. Lower panel: merged image. (F) Deletion of VGluT2 in elPBN neurons by injecting AAV9-hSyn-eGFP-Cre or control AAV9-hSyn-eGFP virus into the right elPBN of VGluT2^fl/fl^ mice. (G) Upper panel: representative coronal slice of VGluT2^fl/fl^ mouse brain depicting the virus injection site in the elPBN. GFP is shown in green. Lower panel: magnified views of the white-boxed region from the upper panel, with transduced cells in the elPBN region indicated by white arrows. Scale bars are 200 µm (low magnification) and 10 µm (high magnification). (H-L) Injury-induced behavioral response: (H) Reflexive responses to different sensory modalities; (I-L) affective-motivational responses to light touch (J), noxious touch (K) and noxious heat (L) in VGluT2^fl/fl^ mice injected with either AAV9-hSyn-eGFP-Cre or AAV9-hSyn-eGFP virus in the right elPBN. Two-way repeated measure ANOVA followed by Sidak’s post-hoc test; *p < 0.05; **p < 0.01; ***p < 0.001; ****p < 0.0001. n = 9 control mice (5 males, 4 females) and 11 Cre mice (4 males, 7 females). Individual mice are represented by scatter points. Triangle symbols indicate female and circle indicate male mice. Ctrl= control virus AAV9-hSyn-eGFP, Cre= AAV9-hSyn-eGFP-Cre virus, ns= non-significant, Inj.= injured paw, Uninj. = uninjured paw. All data are presented as mean ± SEM.

### VGluT2 deletion in elPBN neurons reduces injury-induced pain sensitization

To investigate the role of glutamate neurotransmission within the elPBN in pain-related behaviors, we deleted VGLuT2 in the elPBN using a similar experimental strategy as in the elPBN PACAP deletion. Specifically, elPBN VGLuT2^fl/fl^ mice received stereotaxic injections into the right elPBN with either a Cre-expressing virus (AAV9.hSyna-eGFP-Cre) or a control GFP virus (AAV9.hSyn-eGFP) (**Figure 4F**). As illustrated in **Figure 4G**, virus injection was localized to elPBN neurons. A week after sciatic nerve cuff implantation, we assessed pain sensitivity using nociceptive behavioral tests described in previous experiments. Cre-expressing virus-mediated VGluT2 deletion in elPBN significantly reduced cuff-induced cold (p < 0.0001), tactile (p < 0.0001), pressure (p < 0.0001), and heat (p < 0.001) hypersensitivity in the hind paw ipsilateral to nerve injury compared to control mice (**Figure 4H**). Despite this reduction, mice with VGluT2 deletion in the elPBN retained modest but significant (p < 0.05) injury-induced cold and tactile hypersensitivity, as responses of the paw ipsilateral to the sciatic nerve injury showed a slight but significant increase in sensitivity to cold and tactile stimuli, compared to the uninjured paw. In contrast, injury-induced pressure and heat hypersensitivity were no longer detectable in VGluT2 deletion mice. These combined findings demonstrate that VGluT2 deletion in the elPBN markedly reduces injury-induced hypersensitivity across all sensory modalities tested.

In the hindpaw contralateral to injury, deletion of VGluT2 in the elPBN resulted in a slight but statistically significant (p < 0.05) reduction in evoked tactile responses compared to control group, while responses to cold, heat and pressure stimuli were unaffected by VGluT2 deletion in the elPBN neurons and comparable to control group (Figure 4H).

We also evaluated protective behaviors and escape-seeking responses in mice with VGluT2 deletion in the elPBN neurons following nerve injury (**Figure 4I**). Compared to control mice, Cre-expressing mice exhibited significantly lower affective-motivational responses in response to light touch (p < 0.0001; **Figure 4J**), noxious touch (p < 0.01; **Figure 4K**), and noxious heat stimulation (p < 0.01; **Figure 4L;)** applied on the hind paw ipsilateral to nerve injury. Together, these findings demonstrate that VGluT2-mediated glutamatergic transmission within the elPBN contributes to both reflexive and affective-motivational components of injury-induced pain sensitization across multiple sensory modalities.

### Differential effects of PACAP and VGluT2 deletion in elPBN neurons on inflammation-induced spontaneous pain-related behavior

We previously found that the activation of the PBN to CeA pathway is necessary for inflammation-induced spontaneous pain-related behavior (Torres-Rodriguez et al., 2024). We therefore evaluated the contributions of PACAP and VGluT2 within the elPBN in the formalin model of inflammatory pain by measuring spontaneous pain-like responses following intraplantar injection of formalin (**Figure 5A**). First, we investigated the effect of constitutive knockout of PACAP. Analysis of nociceptive behaviors over time following formalin injection showed a typical biphasic behavioral response in WT mice (**Figure 5B**). In contrast, constitutive PACAP-KO mice exhibited a significant (p < 0.001) overall reduction in the duration of spontaneous nociceptive behaviors (**Figure 5B**). This reduction was driven by significant (p < 0.01) decreased responses during Phase 1 (p < 0.05) and Phase 2 (p < 0.01). (**Figure 5C**). We next separately quantified time spent licking, lifting, or flinching across both phases. During Phase 1, constitutive KO mice showed a significant selective reduction in lifting behavior (p < 0.05), with no alterations in licking or flinching responses (**Supplementary Figure 1B**). During phase 2, PACAP-KO mice displayed significant decreases in both licking (p < 0.01), and lifting (p < 0.05), while flinching behavior remained unaffected (**Supplementary Figure 1B**). Paw thickness, an indirect measure of inflammation-induced edema, was significantly (p < 0.05 and p < 0.01) increased post-formalin injection in both WT and PACAP-KO mice compared to pre-injection measurements (**Figure 5B inset**), indicating that the reductions in spontaneous nociceptive responses are not accompanied by reductions in paw edema. Unlike in the sciatic nerve cuff model, selective PACAP deletion in elPBN neurons failed to recapitulate the constitutive PACAP-KO phenotype, with formalin-induced spontaneous nociceptive responses and paw edema unaffected in elPBN-specific PACAP deleted mice compared to controls (**Figure 5D and 5E; Supplementary Figure 1C; Figure 5D inset**).

**Figure 5:**
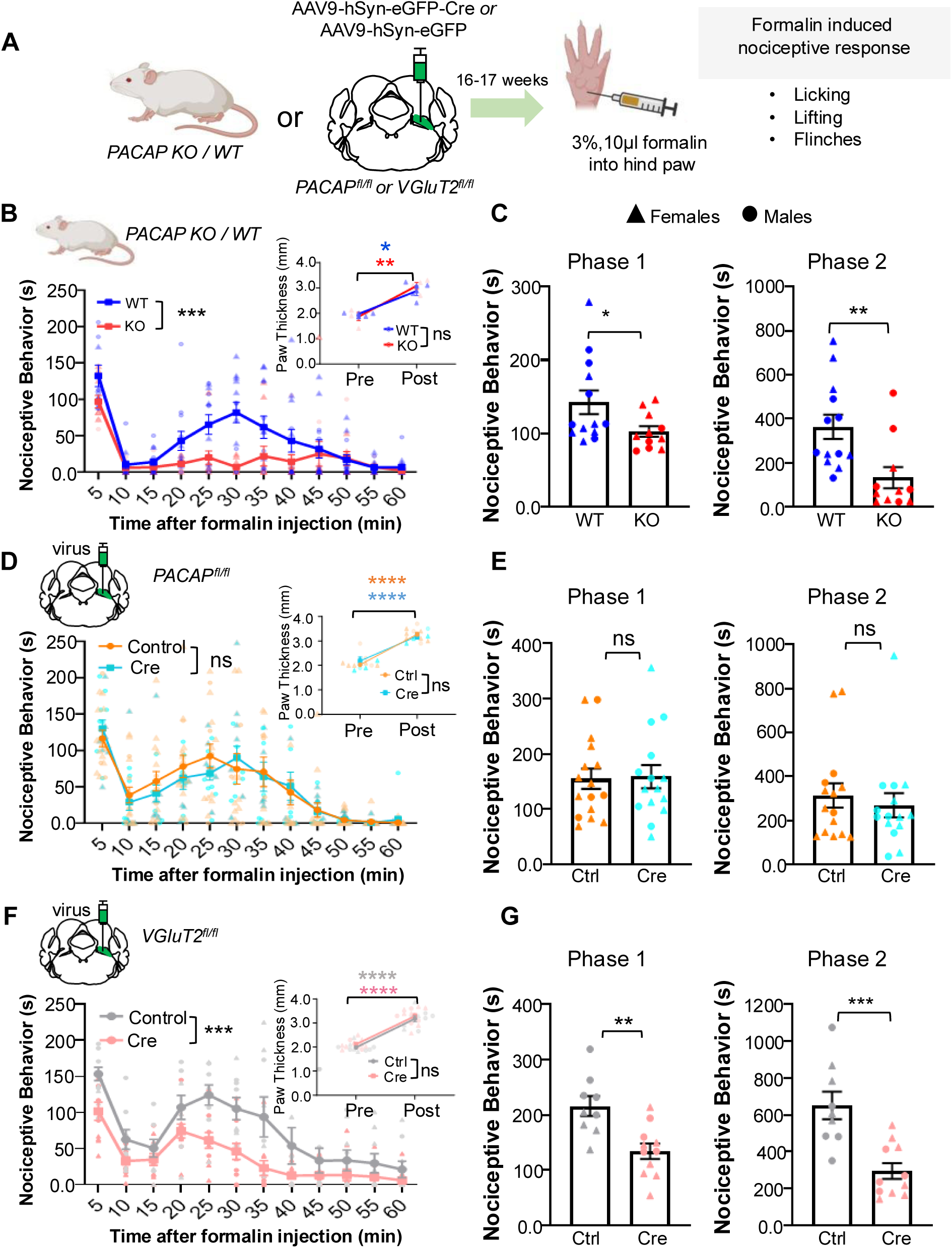
PACAP and VGluT2 deletion in elPBN neurons differently modulate inflammation-induced spontaneous pain-like behavior. (A) Schematic diagram illustrating experimental timeline for measuring formalin-induced spontaneous responses. (B-C) Constitutive PACAP knockout (PACAP-KO) versus wild-type (WT) mice. (D-E) Deletion of PACAP in elPBN neurons (PACAP^fl/fl^ mice injected with AAV9-hSyn-eGFP-Cre or AAV9-hSyn-eGFP virus into the right elPBN). (F-G) Deletion of VGluT2 in elPBN neurons (VGluT2^fl/fl^ mice injected with AAV9-hSyn-eGFP-Cre or AAV9-hSyn-eGFP virus into the right elPBN). (B, D, and F) Time-course of nociceptive behaviors following hind paw formalin injection. Insets: paw thickness pre- and post-injection. (C, E and G) Total time spent in nociceptive behaviors during Phase 1 and Phase 2. Sample sizes: n = 13 WT mice (6 males, 7 females) and 11 constitutive PACAP KO mice (4 male, 7 females); PACAP deletion in elPBN: n = 16 control mice (4 males, 12 females) and 15 Cre mice (8 male, 7 females); VGluT2 deletion in elPBN: n = 9 control mice (5 males, 4 females) and 11 Cre mice (4 males, 7 females). WT= wild-type, KO= knockout, ns= non-significant. Phase 1 is 0–10 minutes and Phase 2 is 10–60 minutes post-formalin injection. Two-way repeated measure ANOVA followed by Sidak’s post-hoc test and Unpaired t-test; *p < 0.05; **p < 0.01; ***p < 0.001; ****p < 0.0001. Scatter points represent individual mice. All data are presented as mean ± SEM. Triangle symbols indicate female and circle indicate male mice.

In contrast, selective deletion of VGluT2 in elPBN neurons significantly reduced total spontaneous nociceptive behaviors (p < 0.001) compared to control virus-infected mice (**Figure 5F and 5G**) without affecting formalin-induced paw edema (**Figure 5F inset)**. Further analysis revealed that elPBN VGluT2 deletion significantly suppressed licking behavior (p < 0.001) during Phase 1, with no effect on lifting or flinching. In phase 2, elPBN VGluT2 deletion significantly decreased both licking (p < 0.0001) and lifting (p < 0.05), whereas flinching remained unchanged **(Supplementary Figure 1D)**. These combined findings indicate that PACAP and glutamatergic signaling within the elPBN contribute differently across pain states, with inflammation-induced spontaneous pain-related behavior engaging elPBN glutamatergic signaling without a detectable requirement for local PACAP.

### Locomotion and affective-like behaviors are largely unaltered following PACAP or VGluT2 deletion in elPBN neurons

Hyperactivity and reduced anxiety-like responses have been reported in mice lacking PACAP (Hashimoto, 2001; Hattori et al., 2012). To determine if selective deletion of PACAP or VGluT2 in elPBN alters locomotor or affect-related behaviors, we performed the open-field test, elevated zero maze, and sucrose splash test **(Supplementary Figure 2A)**. Analysis of locomotor activity in the open field test revealed that the total distance traveled in the whole arena was comparable between control mice and mice with PACAP or VGluT2 deletion. Quantification of time spent in the central zone of the open field, used to assess anxiety-like behaviors, was also comparable between control mice and mice with PACAP or VGluT2 deletion in in elPBN **(Supplementary Figure 2B-E).** Similarly, evaluation of anxiety-like behavior in the elevated zero maze revealed that the percentage of time spent in the open arms and the number of entries into the open arms were indistinguishable in controls and mice with PACAP deletion in the elPBN **(Supplementary Figure 2F and 2G)**. Mice with VGluT2 deletion in the elPBN, however, showed a small but significant (p < 0.05) increase in the time spent in the open arms compared to the control group, while the number of open-arm entries was comparable between groups **(Supplementary Figure 2H and 2I)**. Lastly, neither manipulation altered coat scores and time spent grooming in the sucrose splash test **(Supplementary Figure 2J-M)**. Together, these findings indicate that the effects of PACAP and VGlut2 signaling within the elPBN on pain- and stress-related behaviors occur in the absence of broad alterations in locomotor or affect-related behaviors, although VGluT2 deletion produced a modest change in one anxiety-related measure.

### c-Fos elevation and hypophagia after restraint stress are unaltered in mice with VGluT2 deletion in the elPBN

Deletion of PACAP in elPBN neurons attenuates restraint stress-induced hypophagia and weight loss (Jiang et al., 2023). We therefore asked whether VGluT2-mediated glutamatergic signaling within the elPBN similarly contributes to the hypophagic response to restraint stress. Mice were subjected to a 2-hour acute restraint stress paradigm and body weight was used as an indirect measure of hypophagia (**Figure 6A**). Histological examination and in situ hybridization for *Slc17a6,* the gene encoding VGluT2, revealed a pronounced reduction in *Slc17a6* transcripts within the elPBN of Cre-injected mice relative to controls, confirming effective VGluT2 deletion (**Figure 6B-C**). Acute restraint stress produced robust body weight loss in both control mice and mice with VGluT2 deletion in the elPBN, with comparable magnitude of weight loss between groups (**Figure 6D**). Restraint stress also markedly increased circulating corticosterone levels in both control and Cre groups across both sexes, with no measurable effect of VGluT2 deletion on the corticosterone response under non-restraint or restraint stress conditions (**Figure 6E**). Because PACAP deletion in the elPBN attenuates restraint-induced c-Fos recruitment in the CeA (Jiang et al., 2023), we next asked whether VGluT2 deletion in the elPBN similarly alters CeA recruitment following restraint stress. For this experiment, VGluT2 was unilaterally deleted in the right elPBN. GFP immunoreactivity confirmed viral targeting to the right elPBN and labeled elPBN terminals in the ipsilateral CeA (**Figure 7A-B**). Loss of VGluT2 at elPBN terminals in the CeA was verified by comparing VGluT2 immunoreactivity between ipsilateral (right) and contralateral (left) CeA. Control animals showed comparable VGluT2 labeling bilaterally, whereas Cre-injected mice exhibited significantly reduced VGluT2 immunoreactivity in the ipsilateral CeA (**Figure 7B-C**). Despite marked VGluT2 deletion at elPBN terminals, the number of c-Fos-positive CeA neurons following restraint stress was comparable between Cre-injected and control groups (**Figure 7D**). Together, these findings reveal distinct contributions of PACAP and glutamatergic signaling within the elPBN to acute restraint-stress responses: unlike PACAP loss, VGlut2 deletion did not detectably alter restraint-induced hypophagia or CeA c-Fos recruitment.

**Figure 6.**
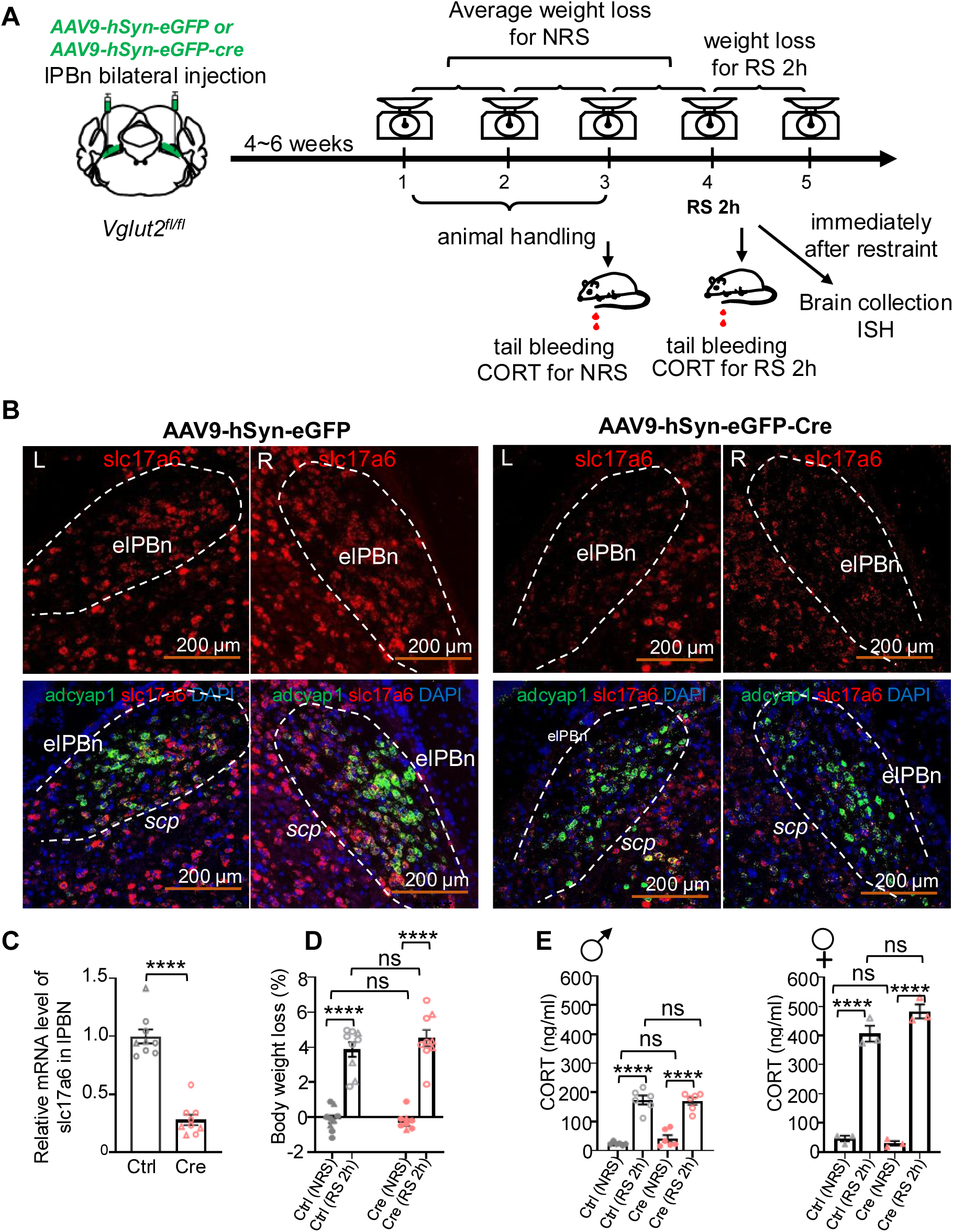
Restraint-induced hypophagia is unaltered after VGluT2 depletion in elPBN neurons. (A) Experimental timeline for VGluT2 deletion in the elPBN and subsequent restraint stress experiments: AAV9-hSyn-eGFP-Cre or the control virus AAV9-hSyn-eGFP was bilaterally injected into the elPBN of VGluT2^fl/fl^ mice. After 4 to 6 weeks, mice underwent 3 days of habituation to handling, followed by 2-hour acute restraint stress on day 4. Tail blood samples were collected to measure corticosterone (CORT) levels under non-restrained (NRS) or restrained (RS) conditions at the 2-hour mark. Body weight was recorded to evaluate hypophagic responses, and brain tissue was harvested for *in situ* hybridization analysis. (B–C) Verification of VGluT2 deletion in elPBN neurons. (B) Representative coronal brain sections using RNAscope *in situ* hybridization showing the elPBN of control virus-injected (left) or Cre-expressing virus-injected (right) mice. VGluT2 (*Slc17a6*) mRNA is shown in red, and PACAP (*Adcyap1*) mRNA is shown in green. Scale bars, 200 µm. (C) Quantification of VGluT2 mRNA signal intensity in the elPBN, demonstrating a marked reduction in VGluT2 expression following Cre-mediated deletion. Quantification of VGluT2 (*Slc17a6*) mRNA signal intensity within elPBN. Values are presented as relative mRNA levels normalized to the mean value of AAV-GFP-injected control mice. (D) Quantification of body weight loss as a proxy for stress-induced hypophagia in mice with VGluT2 deletion in elPBN neurons. (E) Deletion of VGluT2 in the elPBN had no effect on corticosterone increases 2-hour after acute restraint stress. Sample sizes: n = 9 (6 males, 3 females) controls and 9 (6 males, 3 females) Cre mice. All data are presented as mean ± SEM. Scatter points represent individual mice.

**Figure 7.**
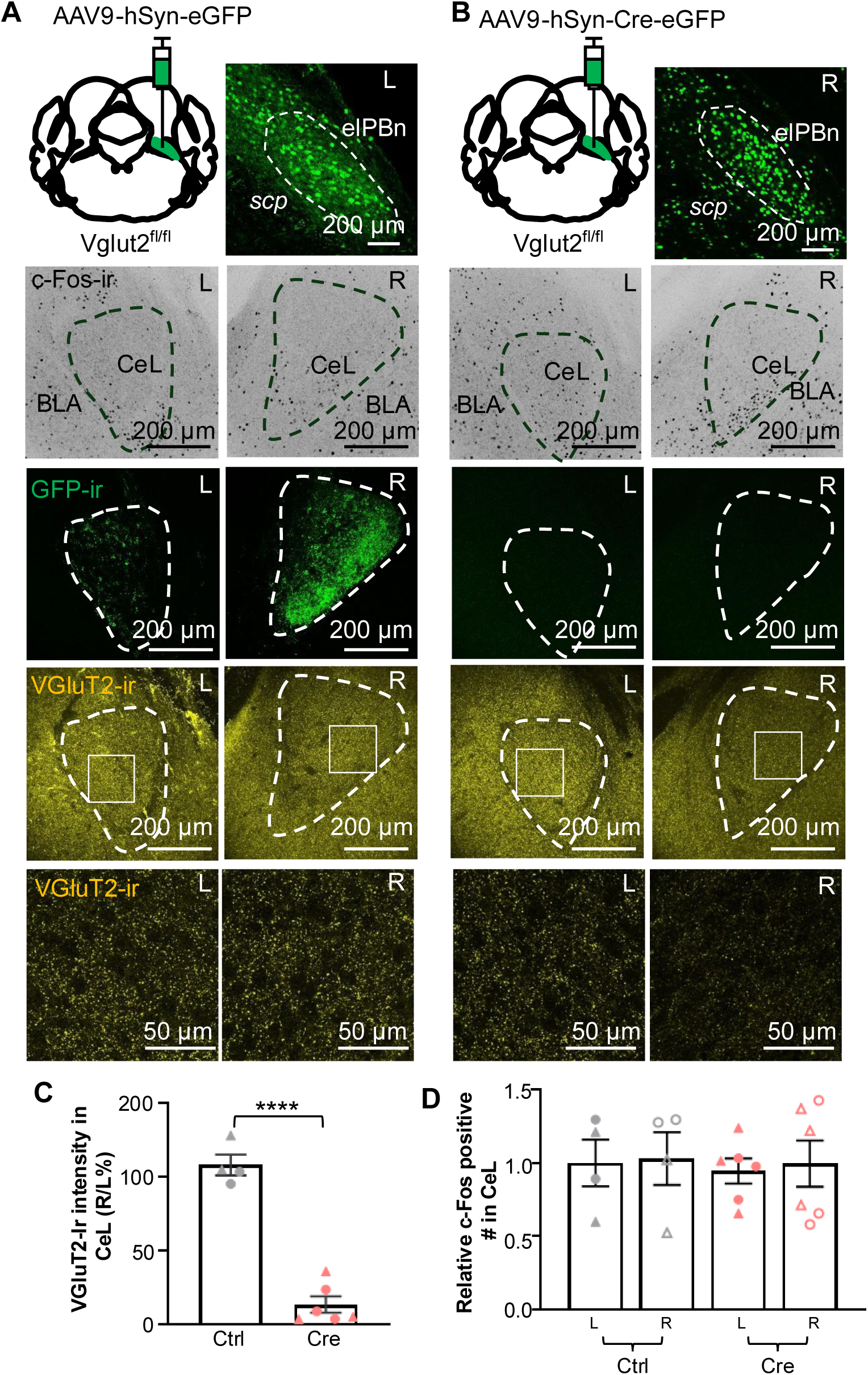
c-Fos expression in the CeA after acute restraint stress is unchanged after deletion of VGluT2 in elPBN neurons. (A-B) AAV9-hSyn-eGFP-Cre (Cre) or the control virus AAV9-hSyn-eGFP (Ctrl) was delivered to the right elPBN in VGluT2^fl/fl^ mice to delete VGluT2 expression. Four to 6 weeks later, c-Fos expression in the CeA was assessed following 2-hours of restraint stress. The sites of viral injection in the elPBn and neuronal terminals in the CeA were marked by GFP immunoreactivity (GFP-ir, shown in green). VGluT2-positive neuronal terminals in the CeA (highlighted in yellow) were identified through VGluT2 immunohistochemistry. (C) Quantification of the intensity of VGluT2-ir signals in the CeA. Results are presented as a percentage ratio of the right (injected) side to the left (uninjected) side (R/L%). (D) The number of c-Fos-positive neurons in the CeA was quantified and normalized to the average count on the left side from mice injected with the control virus. Sample sizes: n=4 (2 males, 2 females) controls and 6 (3 males, 3 females) Cre mice. All data are presented as mean ± SEM. Scatter points represent individual mice.

## DISCUSSION

The generality, and therefore the versatility, of the parabrachial pathway makes it an attractive system for understanding how a relatively small brain region contributes to diverse behavioral responses. Our findings suggest that the neurotransmitter diversity of the elPBN is one important component of this versatility. PACAPergic and glutamatergic signaling make overlapping contributions to nerve injury-induced pain sensitization, but their contributions differ in inflammation-induced spontaneous pain-related behavior and restraint-stress responses (**Figures 1, 2, 4–7**). Thus, the behavioral functions of the elPBN cannot be understood solely from which neurons are recruited or where they project; the signaling molecules used by these neurons also matter. This conclusion is consistent with the growing appreciation that the PBN is heterogeneous at molecular, anatomical, and functional levels and that molecularly defined populations do not necessarily map onto mutually exclusive behavioral functions (Goldstein et al., 2025; Liu et al., 2022; Pauli et al., 2022; Torres-Rodriguez et al., 2024).

The similar effects of PACAP and VGluT2 deletion following nerve injury indicate that both signaling systems make substantial contributions to the sensitized state (**Figures 1, 2, 4**). How these signaling mechanisms are functionally related remains unclear. Their similar effects on nerve injury-induced sensitization, together with previous work, suggest that they may act through parallel mechanisms and may also be released during different phases, perhaps characterized by low- and high-frequency firing rates (Kim et al., 2024), of the pain sensitization process. Extensive electrophysiological work has shown that glutamatergic transmission in the PBN→CeA pathway undergoes marked plasticity during persistent pain (Kato, 2018; Neugebauer V, 2003). Previous pharmacological studies have also implicated PACAP signaling in pain-related plasticity within this circuit (Missig et al., 2017; Missig et al., 2014). Our finding that PACAP loss reduces recruitment of CeA-PKCδ neurons by a normally subthreshold stimulus following nerve injury, while leaving recruitment by strongly noxious stimulation intact (**Figure 3**), is consistent with a greater contribution of PACAP after the pain system has entered a sensitized state. While c-Fos expression does not identify the cellular mechanism underlying this effect, the selective PACAP dependence observed following normally subthreshold stimulation suggests that PACAP becomes particularly important for CeA recruitment in the sensitized state. Mechanistic interactions between peptide and glutamatergic signaling have precedent within the PBN→CeA circuit: calcitonin gene-related peptide (CGRP) facilitates glutamatergic transmission through postsynaptic PKA- and NMDA receptor-dependent mechanisms (Han et al., 2010; Han et al., 2005; Okutsu et al., 2017), and recent work directly measuring endogenous peptide release from PBN terminals demonstrates that CGRP can potentiate glutamatergic signaling in CeA neurons (Lorsung et al., 2025). Whether PACAP similarly modulates glutamatergic transmission remains unknown although PACAP effects on excitability of PKCδ neurons in mouse CeA have been reported (Zhang et al., 2026). While these experiments do not demonstrate PACAP release or peptide–glutamate interactions at these synapses, our finding that PACAP-expressing elPBN terminals provide functional glutamatergic input to CeA-PKCδ neurons (**Figure 4A–E**) identifies an anatomical and functional substrate for such interactions could occur.

The formalin experiments further demonstrate that the similar contributions of PACAP and glutamate following nerve injury cannot be generalized across pain states. VGluT2 deletion in the elPBN reduced spontaneous inflammation-induced pain-related behavior, whereas local PACAP deletion did not (**Figure 5**; **Supplementary Figure 1**). This finding was unexpected based on previous work suggesting a temporal distinction between ongoing nociceptive behavior and the development of plasticity within the PBN→CeA circuit. Sensitization of CeA neurons develops over the first several hours following induction of arthritis (Neugebauer V, 2003), and following formalin injection, CeA ERK activation is detected hours after inflammation but not during the early period of spontaneous pain-related behavior (Carrasquillo & Gereau, 2007). Consistent with this distinction, disruption of CeA mGluR5 signaling or its downstream ERK signaling cascade reduces persistent inflammation-induced hypersensitivity without altering spontaneous formalin behavior (Carrasquillo & Gereau, 2007; Kolber et al., 2010). Similarly, unilateral orofacial formalin produces comparable spontaneous nocifensive behavior regardless of the side of injection, whereas PBN→CeA synaptic potentiation that emerges several hours later is preferentially expressed in the right CeA (Miyazawa et al., 2018). Together, these findings suggest that fast glutamatergic transmission from the elPBN can contribute to the early behavioral response to inflammation, whereas metabotropic signaling and downstream cellular plasticity, including mechanisms potentially influenced by PACAP, may become increasingly important as sensitization develops. The present experiments do not directly test this temporal model; nevertheless, the differential effects of PACAP and VGluT2 deletion across spontaneous inflammatory behavior and nerve injury-induced sensitization are consistent with signaling mechanisms that contribute on different timescales. Determining the temporal dynamics and interaction of PACAP and glutamatergic signaling will be an important next step.

The formalin experiments also reveal an important difference between constitutive and local PACAP loss. Constitutive PACAP knockout reduced formalin-induced spontaneous behavior, whereas adult deletion of PACAP within the elPBN did not (**Figure 5**; **Supplementary Figure 1**). This difference could reflect contributions of PACAP from other sites in the pain neuraxis, developmental consequences of constitutive PACAP loss, or differences in the extent and timing of PACAP deletion. Previous studies have implicated PACAP signaling at spinal and supraspinal levels in inflammatory and neuropathic pain (Mabuchi et al., 2004; Missig et al., 2017; Missig et al., 2014), making contributions outside the elPBN plausible. While the present experiments do not distinguish among these possibilities, the differential effects of constitutive and elPBN-specific PACAP deletion suggest that PACAP signaling outside the elPBN may contribute to inflammatory pain-related behavior. More broadly, the different PACAP and VGluT2 requirements across nerve injury and inflammation (**Figures 1, 2, 4, 5**) indicate that their contributions depend on the pain state and behavioral response being examined and contribute to response versatility within this circuit.

A different relationship between PACAP and glutamatergic signaling emerged during restraint stress. We previously demonstrated that PACAP expression in elPBN neurons contributes to restraint stress-induced hypophagia and recruitment of downstream amygdala circuitry (Jiang et al., 2023). In the present study, VGluT2 deletion did not detectably alter restraint-associated hypophagia, corticosterone responses, or CeA c-Fos recruitment (**Figures 6 and 7**). Thus, the overlapping contributions of PACAP and glutamate to nerve injury-induced sensitization are not required for all elPBN-dependent behaviors. One possible explanation is that different patterns of neuronal activity favor different forms of chemical transmission. Recent studies further demonstrate that peptidergic and glutamatergic signaling from PBN neurons can be differentially engaged. Direct manipulation of neuropeptide release revealed distinct behavioral contributions of peptidergic and glutamatergic transmission in a PBN→CeA circuit (Kim et al., 2024), while direct visualization of LDCV exocytosis from PBN-CGRP terminals demonstrated neuropeptide release during high-frequency stimulation as glutamatergic synaptic responses declined (Lorsung et al., 2025). The different behavioral requirements for PACAP and VGluT2-mediated signaling raise the possibility that transmitter recruitment varies with behavioral state, although PACAP and glutamate release were not directly measured during pain or restraint stress. Differential transmitter release is one possible mechanism, but the findings could also reflect recruitment of different elPBN neuronal populations or projections, differences in downstream targets, or state-dependent interactions between peptide and glutamatergic signaling.

An unexpected finding was the modest increase in open-arm exploration following VGluT2 deletion in the elPBN (**Supplementary Figure 2H–I**). This effect was limited to one measure in the elevated zero maze and was not accompanied by broad changes in locomotor activity or other affect-related measures (**Supplementary Figure 2**); therefore, it should be interpreted cautiously. Nevertheless, it is interesting in light of previous work showing that activation of PACAP-expressing lateral PBN pathways can increase anxiety-related behavior (Missig et al., 2014; Seiglie et al., 2023). The absence of a comparable phenotype following local PACAP deletion (**Supplementary Figure 2F–G**) further emphasizes that manipulating PACAP-expressing neurons, PACAP itself, and glutamatergic transmission from the elPBN are not functionally equivalent. More broadly, this unexpected result raises the possibility that different components of stress- and affect-related behavior depend on different signaling mechanisms within the elPBN.

Determining whether the behavioral effects of PACAP and VGluT2 arise from the same neurons, projections, or synapses will be important for understanding how these signaling systems are functionally related. Direct measurements of PACAP and glutamate release, together with combined manipulation of the two signaling systems, will further help distinguish whether their overlapping contributions to nerve injury-induced sensitization reflect mechanistic convergence, peptide–glutamate interactions, or parallel mechanisms.

Together, our findings show that PACAP and glutamatergic signaling within the elPBN make distinct contributions in mediating pain- and stress-related behaviors (**Figures 1–7**). These results provide functional evidence that the neurotransmitter diversity of the PBN is biologically meaningful and identify the relationship between peptide and glutamatergic signaling as an important question for understanding how this region contributes to diverse behavioral states.

## Acknowledgements

Thanks to Haiying Zhang for mouse colony management and genotyping. This work was supported by the National Institute of Mental Health Intramural Research Program, Project 1ZIAMH002386 to L.E.E, National Center for Complementary and Integrative Health, Intramural Research support to Y.C, and a fellowship from the NIH Center on Compulsive Behaviors to SS. Cartoon diagrams were created with BioRender.com. We thank NIH Section on Instrumentation for helping to design and fabricate the custom-built instruments used in this study, and the NIMH System Neuroscience Imaging Resource (SNIR) and Rodent Behavioral Core (RBC) for expert experimental support.

