## Supplemental figures for "Differential utilization of PACAP and glutamate in parabrachial neurons shapes pain and stress responses"

**Supplementary Figure 1. Time spent in distinct nociceptive behaviors during phase 1 and phase 2 of the formalin test**

(A) Experimental timeline.

(B) Constitutive PACAP-KO mice reduces formalin-induced distinct spontaneous behaviors. n = 13 WT (6 males, 7 females) and 10 constitutive PACAP-KO mice (4 males, 7 females).

(C) PACAP deletion in elPBN neurons does not affect formalin-induced spontaneous behaviors. n = 16 control (4 males, 12 females) and 15 Cre mice (8 males, 7 females).

(D) VGluT2 deletion in elPBN neurons reduces formalin-induced distinct spontaneous behaviors. n = 9 control and 11 Cre mice. ns = non-significant, WT = wild-type, KO = knockout. Unpaired t-test; *p < 0.05; **p < 0.01; ***p < 0.001; ****p < 0.001. Individual mice are represented by scatter points. Triangle symbols indicate female and circle indicate male mice. Phase 1; 0-10 minutes post formalin injection, Phase 2; 10-60 minutes post formalin injection. All data are presented as mean ± SEM.

**Supplementary Figure 2. Locomotion and affect -like behaviors are largely unaltered following PACAP or VGluT2 deletion in elPBN neurons.**

(A) Experimental timeline. Mice underwent a battery of behavioral assays: open field test (OFT, locomotion and anxiety-like behavior); elevated zero maze (EZM, anxiety-like behavior); and sucrose splash test (affect-like behavior).

(B-E) OFT, (F-I) EZM, and (J-M) sucrose splash test. Sample sizes for PACAP deletion in elPBN- OFT: n = 6 control and 3 for Cre mice; EZM test: n = 7 control and 6 Cre mice; splash test: n = 6 for both groups. Sample sizes for VGluT2 deletion in elPBN- OFT, EZM and splash test: n = 9 control and 11 Cre mice. ns= non-significant, Ctrl= control. Individual mice are represented by scatter points. Triangle symbols indicate female and circle indicate male mice. Unpaired t-test; *p < 0.05. All data are presented as mean ± SEM.

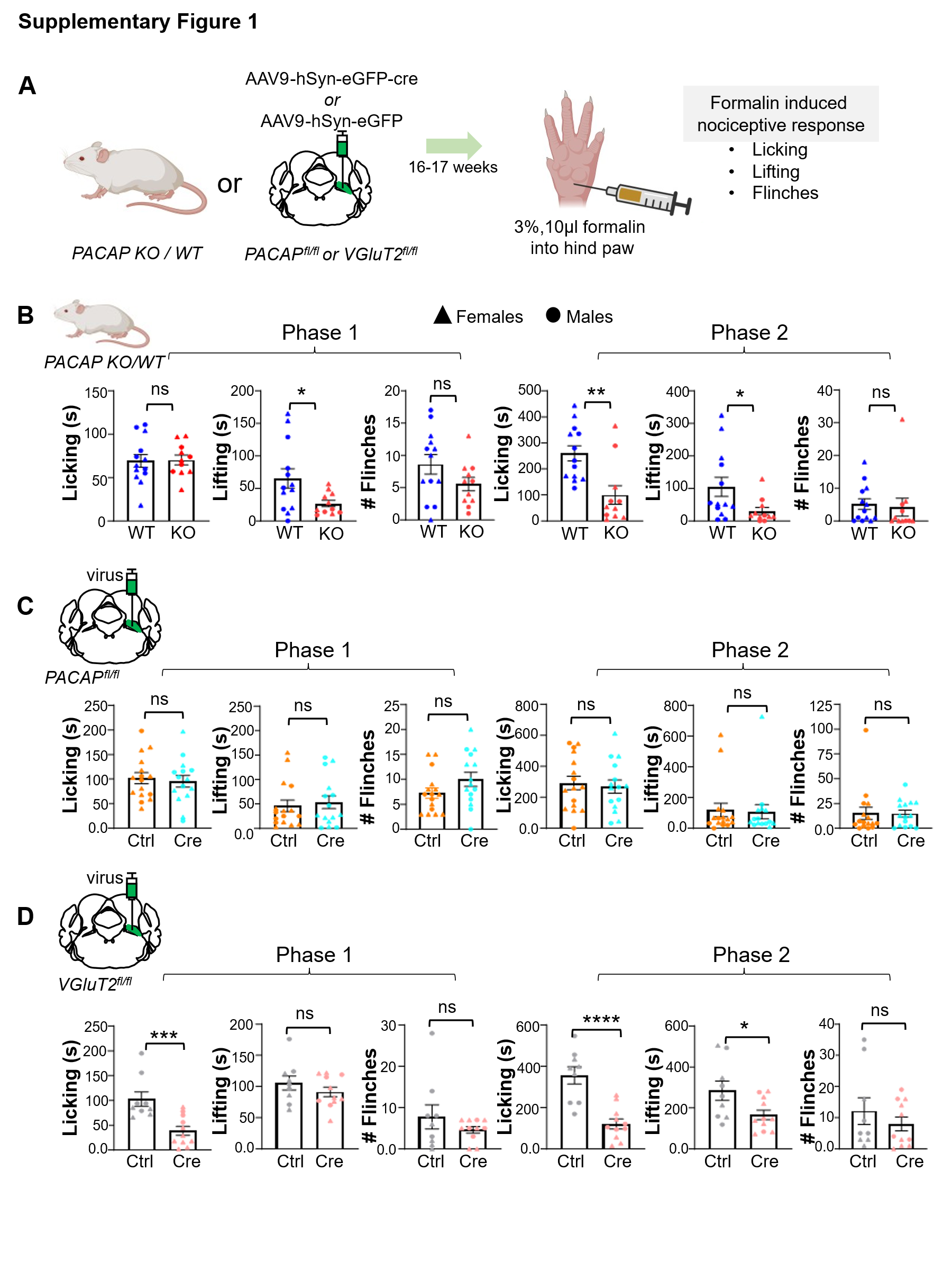

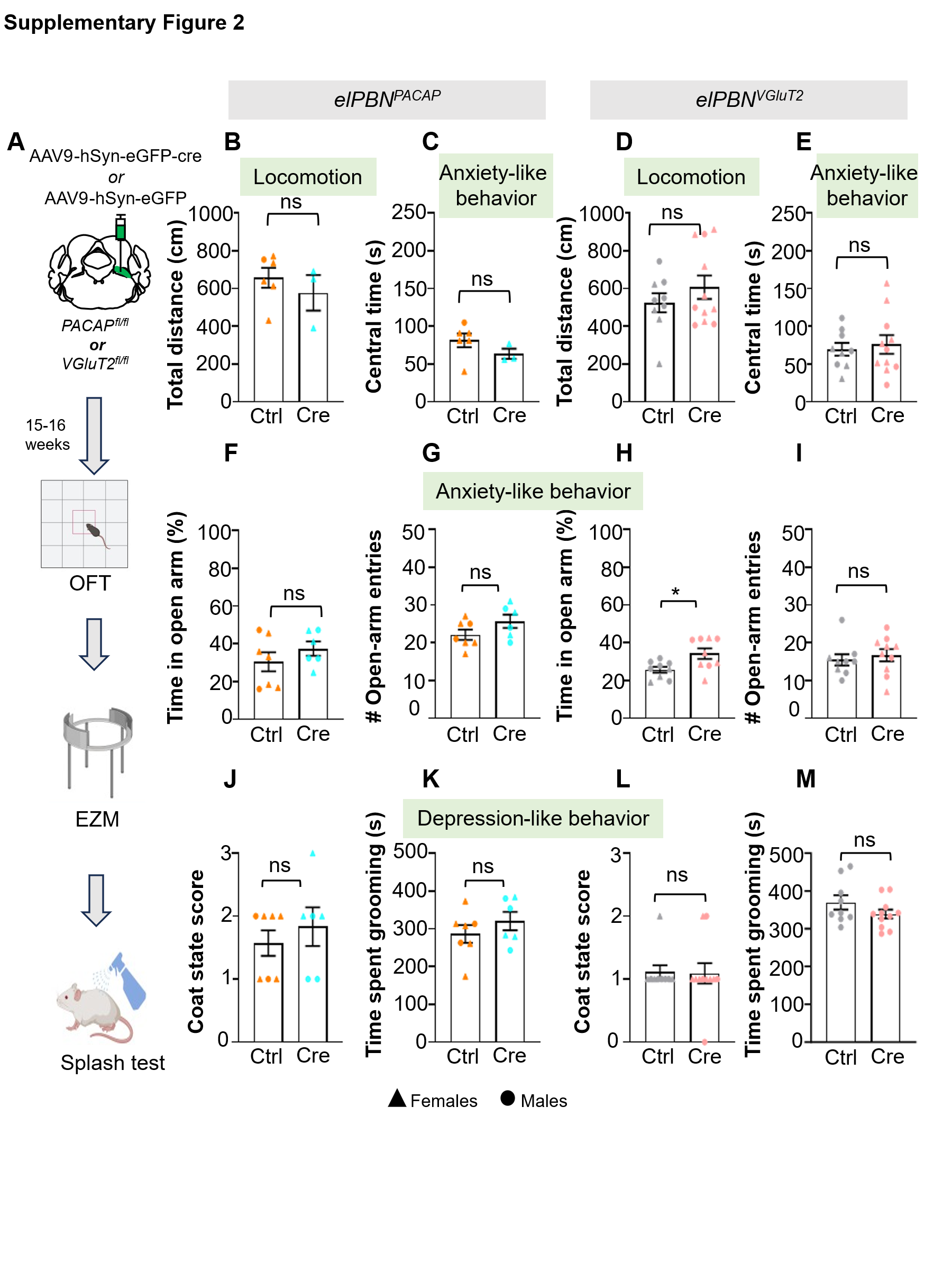
